# An immunoPET probe to SARS-CoV-2 reveals early infection of the male genital tract in rhesus macaques

**DOI:** 10.1101/2022.02.25.481974

**Authors:** Patrick J. Madden, Yanique Thomas, Robert V. Blair, Sadia Samer, Mark Doyle, Cecily C. Midkiff, Lara A. Doyle-Meyers, Mark E. Becker, Muhammad S. Arif, Michael D. McRaven, Lacy M. Simons, Ann M. Carias, Elena Martinelli, Ramon Lorenzo-Redondo, Judd F Hultquist, Francois J. Villinger, Ronald S. Veazey, Thomas J. Hope

## Abstract

The systemic nature of SARS-CoV-2 infection is highly recognized, but poorly characterized. A non-invasive and unbiased method is needed to clarify whole body spatiotemporal dynamics of SARS-CoV-2 infection after transmission. We recently developed a probe based on the anti-SARS-CoV-2 spike antibody CR3022 to study SARS-CoV-2 pathogenesis *in vivo*. Herein, we describe its use in immunoPET to investigate SARS-CoV-2 infection of three rhesus macaques. Using PET/CT imaging of macaques at different times post-SARS-CoV-2 inoculation, we track the ^64^Cu-labelled CR3022-F(ab’)2 probe targeting the spike protein of SARS-CoV-2 to study the dynamics of infection within the respiratory tract and uncover novel sites of infection. Using this method, we uncovered differences in lung pathology between infection with the WA1 isolate and the delta variant, which were readily corroborated through computed tomography scans. The ^64^Cu-CR3022-probe also demonstrated dynamic changes occurring between 1- and 2-weeks post-infection. Remarkably, a robust signal was seen in the male genital tract (MGT) of all three animals studied. Infection of the MGT was validated by immunofluorescence imaging of infected cells in the testicular and penile tissue and severe pathology was observed in the testes of one animal at 2-weeks post-infection. The results presented here underscore the utility of using immunoPET to study the dynamics of SARS-CoV-2 infection to understand its pathogenicity and discover new anatomical sites of viral replication. We provide direct evidence for SARS-CoV-2 infection of the MGT in rhesus macaques revealing the possible pathologic outcomes of viral replication at these sites.

**Graphic Abstract:** PET/CT detected SARS-CoV-2 infection of 4 different tissues in the male genital tract illuminates the cause of COVID-19 clinical sequalae of male sexual health and fertility

**Figure 1.**
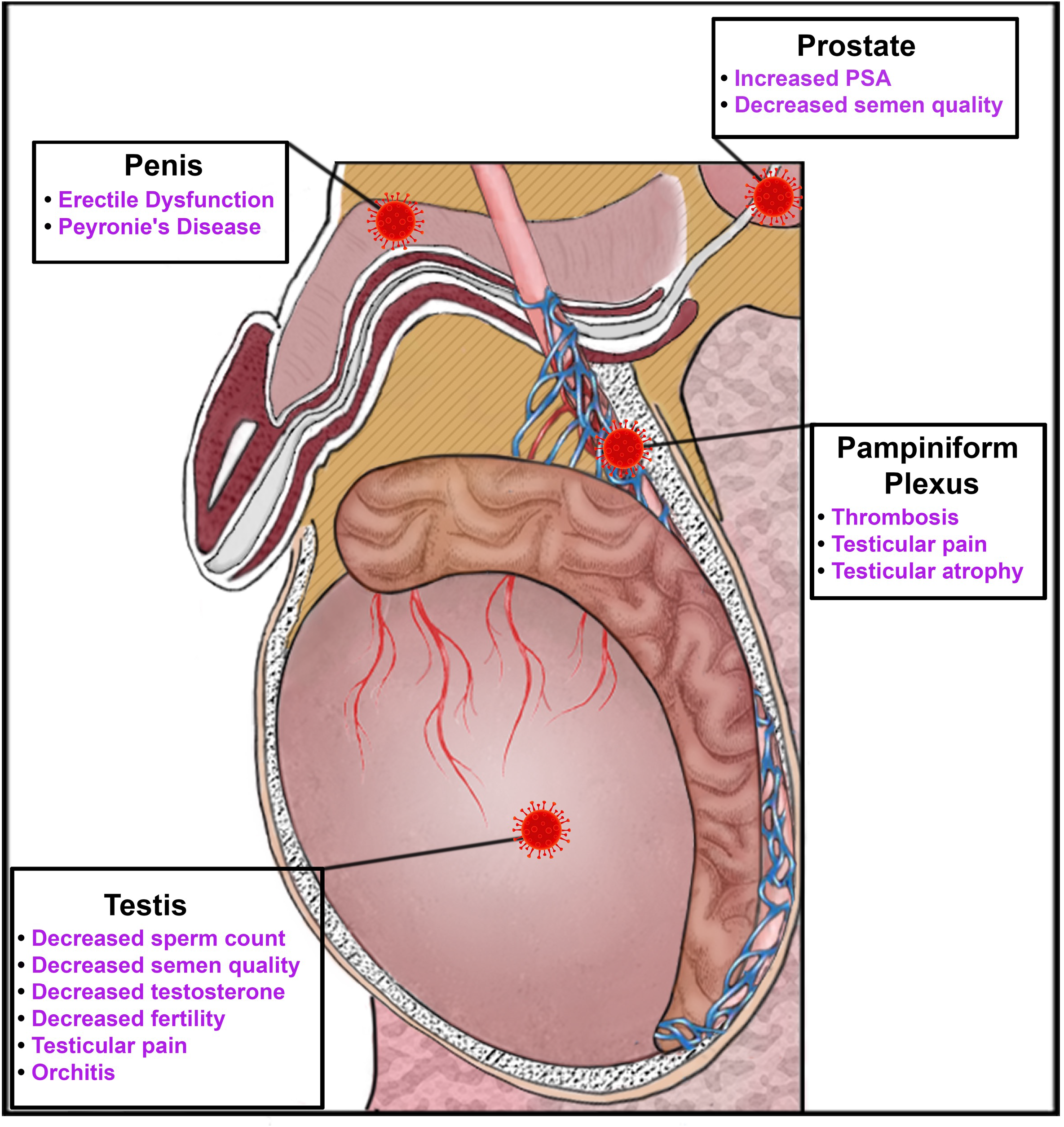
Diagram shows schematic illustration of the male genital tract of the rhesus macaque. Virus icon shows sites of SARS-CoV-2 PET signal. Text highlighting the clinical sequalae associated with each sight of infection is shown in text adjacent to each infection site.

## Introduction

The COVID-19 pandemic has exposed the broad systemic impact that can be caused by infection with a respiratory virus. Disease associated with SARS-CoV-2 infection starts with respiratory pathologies and subsequently can extend to other organ systems. There is now ample evidence that SARS-CoV-2 can disseminate and replicate in tissues beyond the respiratory tract. A clear example is infection in the gastrointestinal (GI) tract^1^. GI symptoms, including diarrhea, have been reported by individuals with mild COVID-19, and hospitalized patients have exhibited more severe symptoms such as ischemia and GI bleeds ^2–4^. In addition, it is now well established that virus is shed through the GI tract in most infected individuals and wastewater screening has become an important tool for disease surveillance^5, 6^. Although less studied, many other tissues have been found to harbor SARS-CoV-2. Multiple groups have shown the presence of viral RNA in cardiac, renal, and brain tissues^7–10^. There is also some evidence of virus in the male genital tract (MGT) ^11, 12^. Furthermore, symptoms associated with all these organ systems have been regularly reported^13^. Likewise, other RNA viruses have documented early dissemination to distal tissues that manifest infection-related pathology over the long term, including (but not limited to) polio, mumps, Ebola virus, Zika virus, and SARS-CoV-1^14, 15^. For example, studies of autopsy tissue from fatalities of SARS-CoV-1 suggested that it causes inflammation of the testes (orchitis)^15^.

However, it is not yet clear if the diverse pathologies associated with SARS-CoV-2 infection are due to secondary effects of systemic inflammation or direct infection of tissues at these distal sites. Human studies have relied on biopsies and autopsy samples to investigate viral replication in sites other than the respiratory tract. The biopsy/autopsy samples only offer a snapshot of the *in vivo* dynamics of disease, and there are ethical and technical difficulties in obtaining them. Using animal models of infection allows for a more thorough collection and investigation of affected tissues. To gain critical insights into systemic infection by SARS-CoV-2, new animal models are needed to determine the extent of disseminated infection and its relationship to pathogenesis.

SARS-CoV-2 infection in the non-human primate (NHP), rhesus macaques (*Macaca mulatta*), recapitulates mild to moderate human disease^16–18^. Infected macaques exhibit viral shedding through the respiratory tract and viral pneumonia similar to the mild form seen in humans. The architecture of the respiratory tract is also generally conserved between humans and macaques making them an ideal model for studying SARS-CoV-2 infection. Infection of other organ systems has also been observed in macaques, most commonly in the GI tract^19, 20^. To gain insights into the spatiotemporal dynamics of SARS-CoV-2 during infection in the rhesus macaque model, we adapted an immunoPET methodology that we are currently using to study various aspects of simian immunodeficiency virus (SIV) acquisition and pathogenesis^21^. ImmunoPET is a molecular imaging technique that combines the specificity of an antibody-based probe labeled with a radioisotope with the *in vivo* imaging power of combined positron emission tomography-computed tomography (PET/CT). ImmunoPET was originally developed and has been widely used in cancer research. Recently, with the advent of new antibodies and better radioisotopes, immunoPET has been extended to studying many other biological processes, including the *in vivo* dynamics of pathogens ^22, 23^. ImmunoPET allows for repeated and specific imaging of virally infected cells *in vivo* by using a radioisotope labeled antibody targeting a viral protein. The non-invasive nature of PET/CT imaging allows for unbiased discovery of novel tissue sites of infection through whole body imaging. Furthermore, the cell-associated PET signal persists in tissue allowing for a radioactive probe-guided necropsy to help determine the precise location of infected cells.

We have previously reported the early development of an *in vivo* antibody-based probe against SARS-CoV-2 utilizing fluorescent tagging of the F(ab’)2 of the anti-spike IgG CR3022^24^. CR3022 was one of the first monoclonal antibodies identified that bound tightly to SARS-CoV-2. It was originally derived from an individual infected with SARS-CoV-1 but also exhibits tight binding to the spike protein of SARS-CoV-2. Here, we extend the use of the F(ab’)2 of the anti-spike IgG CR3022 labelled with copper 64 (Cu^64^) for immunoPET and targeted necropsy to study systemic SARS-CoV-2 infection in the rhesus macaque model. Our results show the utility of this approach in investigating SARS-CoV-2 pathogenesis in the respiratory tract and in uncovering novel anatomical sites of infection. Interestingly, we detected a robust and dynamic signal in the MGT including the prostate, penis, and testicles. This observation is consistent with emerging and ongoing clinical observations of orchitis, oligo-/azoospermia, and erectile dysfunction, and reveals these comorbidities are likely a consequence of the direct viral infection of the tissues of the MGT. The successful development of an immunoPET probe to study SARS-CoV-2 in the rhesus macaque challenge model will allow longitudinal studies to gain insights into SARS-CoV-2 progression, dissemination, and the development of comorbidities.

## Results

### Description of macaque studies and PET/CT-guided necropsy

The basic process and workflow of the PET/CT guided necropsy method is shown in Fig 1A. The PET/CT guided necropsy approach consists of three separate PET/CT scans that are used to map probe signal at the whole animal, organ, and tissue levels. The first scan is typically ∼16-24 hours after the injection of the radio labelled F(ab’)2 probe allowing for movement into the tissues^21^. This whole-body PET/CT scan (Scan 1) identifies “hot” organs and tissue areas. These tissue areas are collected at necropsy immediately following the scan and subjected to a second PET/CT scan (Scan 2). Tissues containing probe signal are cut into small blocks, placed in cryomolds, and then rescanned (Scan 3) to identify individual “hot” tissues/blocks that likely contain foci of virally infected cells. These “hot” tissues can then be used for downstream characterization including RNA quantification and different types of microscopic analyses characterizing virally infected cells.

**Figure 1.**
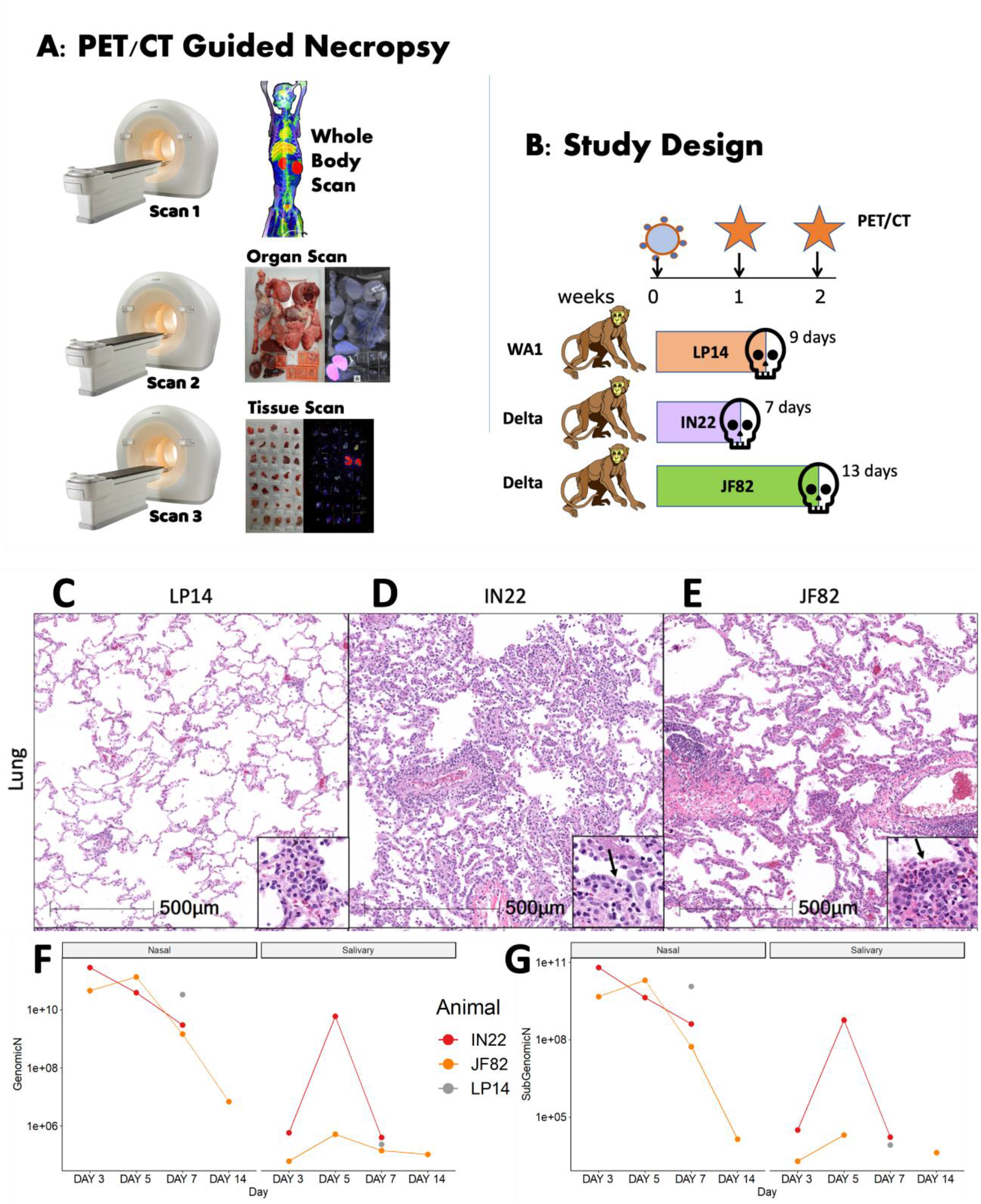
Study design and viral analysis of infected macaques. (A) Workflow of PET/CT guided necropsy. (B) Schematic showing the study design of probe administration, PET/CT scans, and infection. (C-E) Lung lesions were consistent with prior findings in NHPs infected with SARS-CoV-2 and varied from minimal in LP14 (C), moderate in IN22 (D), and mild in JF82 (E). Insets demonstrate the inflammatory infiltrate and in IN22 and JF82, type II pneumocyte hyperplasia (arrows). (F and G) Viral load measurements for all three animals. (F) shows copies/swab of genomic N while (G) shows copies/swab of subgenomic N.

The SARS-CoV-2 pilot infection study design with 3 male rhesus macaques is shown in Fig 1B. Based on our previous studies utilizing a fluorescently tagged F(ab’)2 probe, we decided to perform the first PET/CT scan 1 week after challenge. In the first study, we infected a male rhesus macaque (LP14) with the WA1 isolate of SARS-CoV-2 and performed a single PET/CT scan followed immediately by necropsy. In the second study, two male rhesus macaques were infected with the Delta variant of SARS-CoV-2, and both underwent a PET/CT scan 1 week after infection. One of the animals (IN22) was scanned twice after a single probe injection and necropsied after the 2^nd^ scan. The early scan of IN22 was done 3 hours after injection of the ^64^Cu-labelled F(ab’)2 probe and a second whole-body scan was done ∼21 hours after probe injection with the PET/CT guided necropsy done immediately after. For the third animal (JF82), the 1^st^ PET/CT scan at week 1 post infection was performed ∼22 hours after IV probe administration. On day 13, this same animal, JF82, was evaluated at week 2 post-infection by whole-body scan 18 hours after probe administration followed by necropsy and subsequent scans.

### SARS-CoV-2 infection characteristics

SARS-CoV-2 infection of the 3 animals was monitored via classical hematoxylin and eosin (H&E) staining for evaluation of lesions in lung tissue (Fig 1C-E) and via quantification of genomic and subgenomic RNA in nasal swabs and saliva (Fig 1F, G). Pathology varied from minimal in LP14 (Fig 1C), to mild in IN22 (Fig 1D) and JF82 (Fig 1E). These findings are consistent with the pulmonary pathology of SARS-CoV-2 infected rhesus macaques observed in other studies performed at the Tulane National Primate Research Center^17, 25^. The pulmonary pathology consisted of variable interstitial inflammation and type II pneumocyte hyperplasia (IN22 and JF82), which in the most severely affected animal (IN22) also exhibited prominent atypia. Fig 1F and G shows the level of genomic (Fig 1F) and subgenomic (Fig 1G) RNA in nasal swabs and saliva for all 3 animals. No obvious differences were noted in viral loads between the 3 animals.

### Distribution of SARS-CoV-2 signal after 1 week of infection with the WA1 isolate

The three PET/CT scans of LP14, including the whole body (Fig 2A and B, S1-video), the organ scan (Fig 2C, D, and E, S2-video) and the tissue scan (Fig 2F), revealed probe signal at a number of distinct anatomical sites. As expected, there was a strong signal in the kidneys, which is a consequence of the excretion of the radiolabeled probe. To better evaluate the probe signal in the lungs, the 3D reconstruction of the lungs was isolated from the PET datasets, and the lung signal was projected over the CT reconstruction of the skeleton (Fig 2G and H, S3-video). The signal associated with the lungs was diffuse and quite uniform throughout the tissue as observed in the whole animal scan. This is more evident in the projection of the lung signal without CT (Fig 2I and J, S3-video). However, the lung signal was more evident in the post necropsy organ scan when the intact respiratory tract including the tongue were scanned again (Fig 2C). In this case, lungs were no longer inflated within the body increasing signal density, leading to higher signal in the 2^nd^ scan. Two different respiratory tract z-series images (Fig 2K and L) reveal a focus of signal associated with the base of the tongue, overlapping with the pharynx as evidenced by the cartilaginous structures seen in the CT. There were abundant small foci of signal distributed throughout the lung tissue, often adjacent to the trachea and bronchi as they branch out into the lungs. In addition, the z-series’ reveal that the observed PET signal is associated with the tissues and not with the open airways of the bronchus and ever narrowing bronchi. Immunofluorescence microscopy revealed areas of infected cells lining alveoli in lung tissue sections that had PET/CT signal (Fig 2M and N) confirming the specificity of the PET signal.

**Figure 2.**
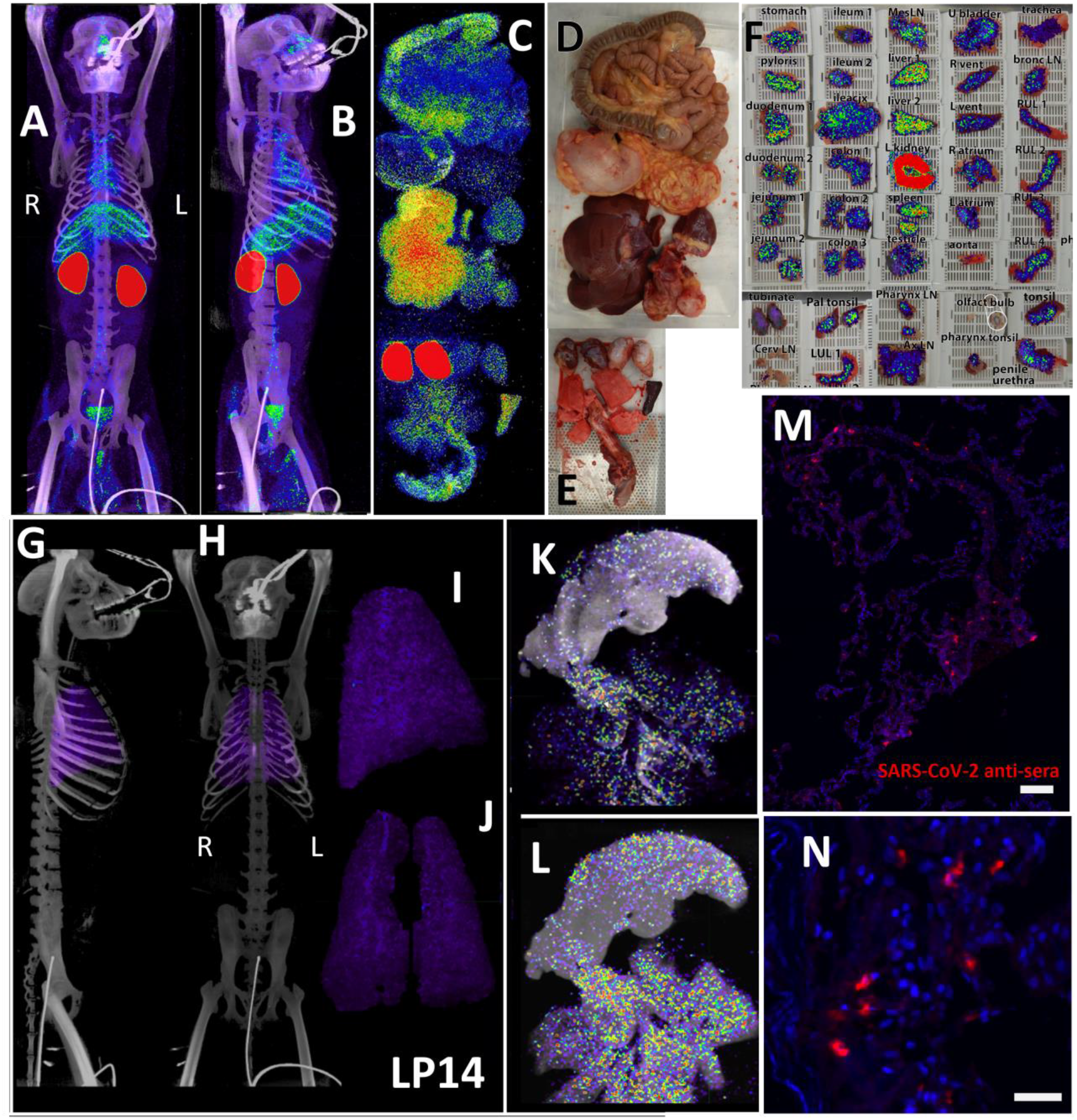
LP14 PET/CT guided necropsy. (A and B) Whole-body PET/CT scans of LP14 8 days post-infection. Front view (A) and rotated 45° (B) both shown. PET signal is display as SUV. (C) Post necropsy PET/CT organ scan of LP14. (D and E) Images of tissues scanned in (C). (F) Overlay of PET signal onto photograph of small pieces of tissue in cassettes. (G and H) Lung PET signal was isolated and overlaid on whole body CT scans. Side (G) and front (H) view. (I and J) Isolated lung PET volumes used in G and H are shown independently, side view (I) and front view (J). (K and L) Single axial z-slice images of respiratory tract PET/CT signals are shown, each image represents a single z-plane from scan shown in C. (M and N) Fluorescent microscopy images of LP14 lung tissue blocks. Red is SARS-CoV-2 anti-sera and blue is Hoechst nuclear stain. Scale bars 100 µM (M) and 25 µM (N).

In contrast to the weak, diffuse signal associated with the lungs, there was a strong and more focal PET signal within the MGT as shown in the front (Fig 3A) and side (Fig 3B-C) views (S4-video). The testes have a generally diffuse signal with some concentration of signal at the dorsal and ventral surface of both LP14 testes. The MGT of the rhesus macaque is similar to that of humans but is distinct in several ways. Most notably, it is primarily retracted within the body covered by a prepuce with a small bone located within the glans penis known as the baculum (visible in the CT scan of the animals, Fig 3B and C, red arrow). There was a very pronounced PET signal associated with the root of the penile tissues, which is buried within the abdomen. The 3D PET signal was projected in green in a front (Fig 3D) and side view (Fig 3E) to better illustrate the signal associated with the MGT and to visualize the signal from the root of the penis (see S5-video). The localization of the PET signal for the penis and testes in the context of the skeletal CT signal is shown in three different projections in panels 3F, 3H, and 3J, and isolated in panels 3G, 3I, and 3K (S6-video). The testis overlays shown in Fig 3L-O reveal that the PET signal is primarily associated with the testis and is distributed throughout the testicular tissue. There is also some potential signal associated with the epididymis.

**Figure 3.**
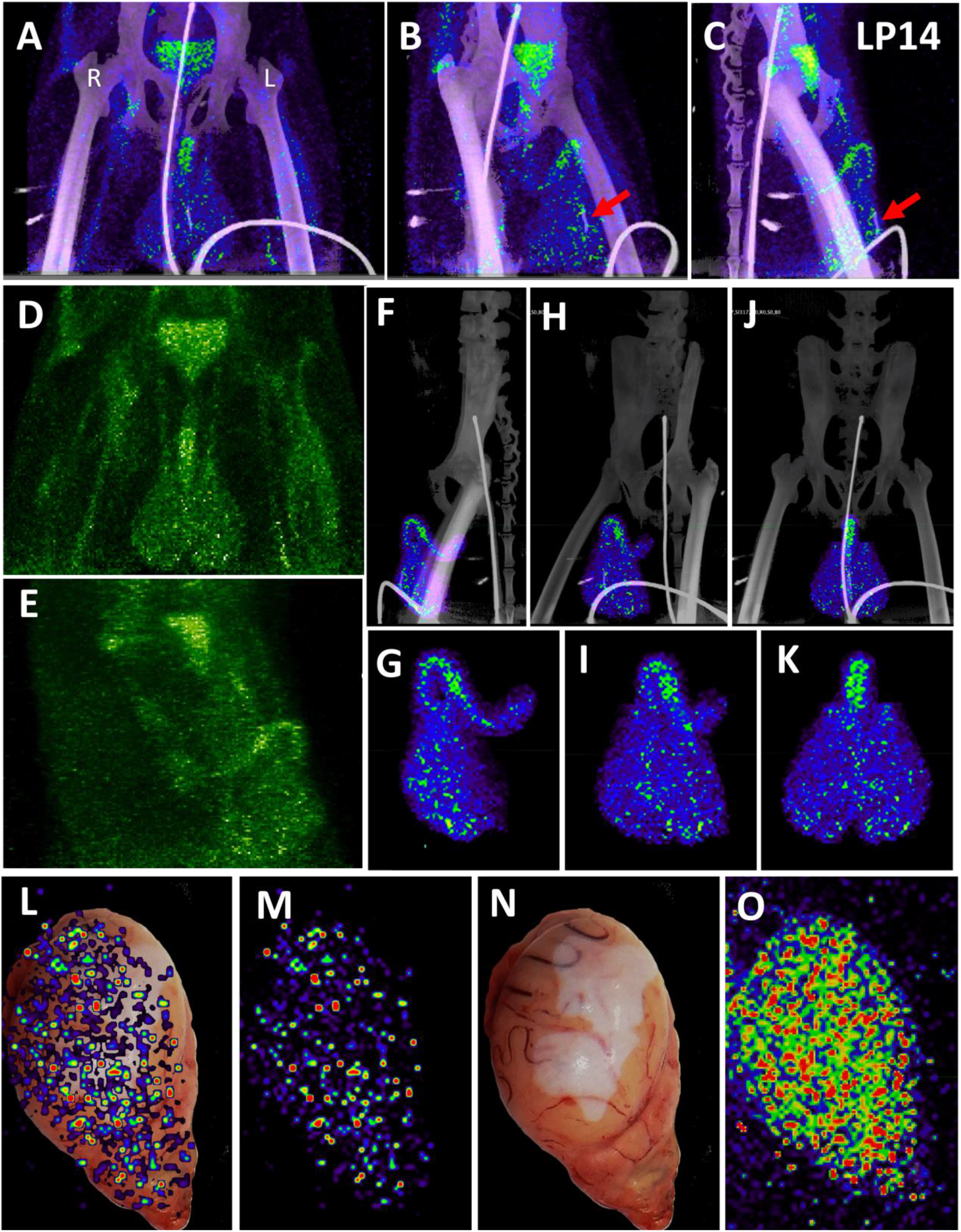
Male genital tract signal in LP14. (A, B, and C) PET/CT images highlighting the lower abdomen of LP14 from the whole-body scan. Front (A), rotated 45° (B), and side (C) views are all shown. Right and left labeled in front view. Red arrow in B and C shows location of baculum in CT scan. (D and E) PET signal with CT overlay removed to highlight signal in MGT, front (D) and side (E) views shown. (F, H, and J) Isolated 3D volume of MGT from whole body scan overlaid with CT images. Side (F), rotated 45° (H), and front (J) views shown. (G, I, and K) Isolated 3D volume of MGT used in overlays (F, H, and J) shown with same views. (L) Overlay of single z-plane of PET signal from organ scan onto image of testis of LP14. (M) PET signal from single z-plane of organ scan used in (L). (N) Image of LP14 testis used in (L). (O) 3D volume of PET signal from organ scan of single testis in previous panels.

In anticipation of a potential signal in the testes, which are known to express the SARS-CoV-2 receptor ACE2, we had designated the testes to be a component of the organ scan post necropsy. Unfortunately, much of the penis tissue was discarded at the necropsy of LP14, though we were able to isolate a small piece of penile tissue remaining within the abdomen. This small piece of penile tissue retained PET signal as can be seen in Fig 2 (Fig 2F: row 8 column 5, Fig S2). To potentially identify infected cells within the testicular tissue of LP14, we evaluated tissue sections for expression of SARS-CoV-2 proteins and host proteins using immunofluorescence microscopy. In LP14, we found that ACE2 is expressed in the peritubular myoid and Sertoli cells surrounding/lining the base of the seminiferous tubules (Fig 4A). Next, we stained testicular tissue sections from LP14 with an anti-SARS-CoV-2 guinea pig serum. Infected cells were evident in seminiferous tubules but were not widely distributed and were instead localized in small clusters of cells at the base of the seminiferous tubules (Fig 4B). Infection of Leydig and other interstitial cells could not be evaluated due to loss of these cells during tissue processing. To further investigate the phenotype of infected cells, we stained with markers to help identify peritubular myoid cells (smooth muscle actin) and Sertoli cells (vimentin). SARS-CoV-2 anti-serum colocalized with vimentin staining at the base of the tubules, indicating that Sertoli cells are being infected (Fig 4C). However, SARS-CoV-2 staining was sometimes observed in cells negative for vimentin indicating that cellular targets exist in addition to Sertoli cells (presumably germ cells because of their localization) (Fig 4C, arrow heads).

**Figure 4.**
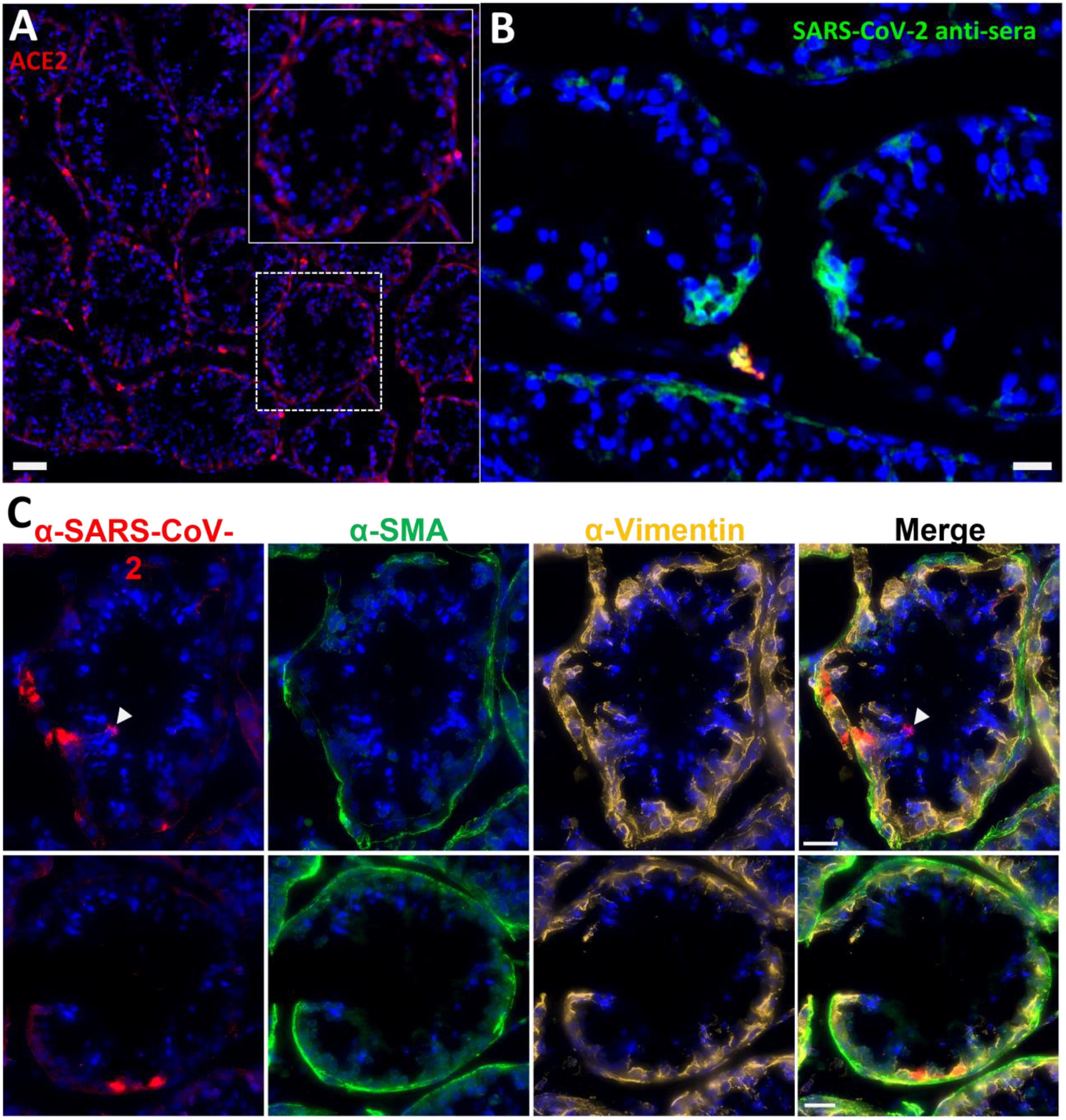
Immunofluorescence of LP14 Testes. (A) Fluorescent microscopy image of LP14 seminiferous tubules. ACE2 staining shown in red, Hoechst nuclear staining shown in blue. Inset shows zoom in of single tubule to better view ACE2 staining in Sertoli and myoid cells. Scale bars 50 µM. (B) Fluorescent microscopy image of LP14 testis shows infected cells. SARS-CoV-2 anti-sera staining in green, background fluorescent in red, and Hoechst nuclear staining in blue. Scale bars 25 µM. (C) Microscopy images of two tubules containing infected cells (top and bottom rows). Red is SARS-CoV-2 anti-sera, green is smooth muscle actin, gold is vimentin, and blue is Hoechst nuclear stain. Possible infected germ cells marked by arrow heads. Scale bars 20 µM.

### Distribution of SARS-CoV-2 Delta variant signal after 1 week of infection

Because of the mild disease manifestations of SARS-CoV-2 WA1 seen in many rhesus macaque studies, we utilized the Delta variant for the next set of animals, which had become the dominant circulating variant at the time of this study^26^. For IN22, at 1-week post infection, we performed 2 PET/CT scans at 3- and 21-hours post probe injection to gain insights into the dynamics of the probe distribution over time. This is an important aspect of probe function because as the probe distributes from circulation into the tissue it will encounter virally infected cells which will affect its distribution pattern. The three PET/CT scans of IN22, the whole body (Fig 5A-D and S7-video, S8-video), the organ scan (Fig 5E-H and S9-video) and the tissue scan (Fig 5I) are shown in Fig 5.

**Figure 5.**
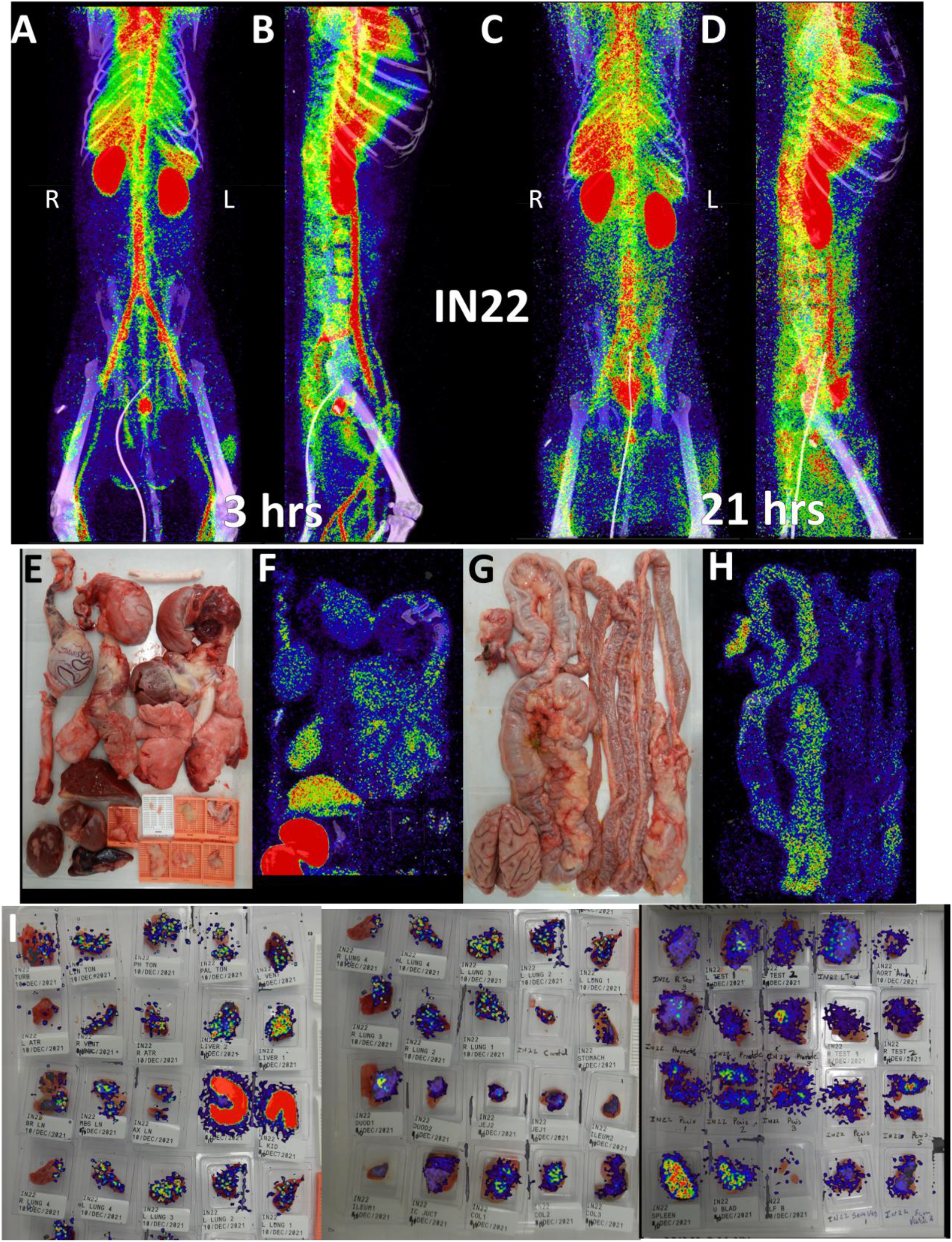
IN22 PET/CT guided necropsy images. (A and B) Whole-body PET/CT scans of IN22 obtained 3-hours after probe administration. Front view (A) and side (B) both shown. (C and D) Whole-body PET/CT scans of IN22 obtained 21-hours after probe administration. Front view (C) and side (D) both shown. Right and left labeled in front views (A and C). (E) Organ tray post necropsy and (F) PET/CT image. (G) Second organ tray post necropsy and (H) PET/CT image. (I) Overlay of PET signal onto photograph of tissue cassettes.

The front and side views of the 3-hour scan (Fig 5A, B and S7-video) and 21-hour scan (Fig 5C, D and S8-video) reveal a highly dynamic system of probe distribution demonstrating that the timing of the PET/CT scan after probe injection is an important consideration. The 3-hour scan clearly captures a robust probe signal in the vasculature and chambers of the heart, consistent with a large fraction of the probe still circulating in the blood after IV injection. A strong lung signal is apparent in the 3-hour scan in contrast to the previous animal. Some signal was also observed in the MGT with a very strong signal observed in the prostate in the 3-hour scan. In the 21-hour scan, the lung probe signal remained much greater than observed in LP14 and was further distributed throughout the tissue. The prostate signal remained but decreased substantially in the 21-hour scan while signal in the MGT was further amplified. For the most part, an increase in signal was seen at tissue sites of probe labeling in the 21-hour scan. In contrast, the vasculature signal decreased substantially, consistent with the movement of the probe from blood, into tissues. However, some discrete probe labeling of certain vascular sites remained, possibly indicating infection of the vasculature.

To better evaluate the probe signal in the lungs after Delta infection, the 3D reconstruction of the IN22 lungs was isolated from the PET datasets, and the lung signal was projected over the CT reconstruction of the skeleton (Fig 6A-D, S10-video). Both the 3-hour and 21-hour signals are significant and localized. The lung rotation series (Fig 6E and F) reveal a major signal associated with the caudodorsal portion of the right lung and less signal associated with the left lung. Single z images of the PET and CT signal in sagittal and transverse sections is shown in Figs 6G-J to better facilitate the analysis of the relationship of CT revealed lung pathology with the PET signal. This comparison reveals an overlap of the opaque lung signal and greater PET signal in the right lung in contrast to the more CT transparent left lung tissue (Fig 6G-J). The stronger PET signal in the right lung observed in the whole-body scan is recapitulated in the organ scan (Fig 6K-O). This animal had overt gross pathology associated with the dorsal aspects of the right lower lung lobe as highlighted in the magnified inset (Fig 6L).

**Figure 6.**
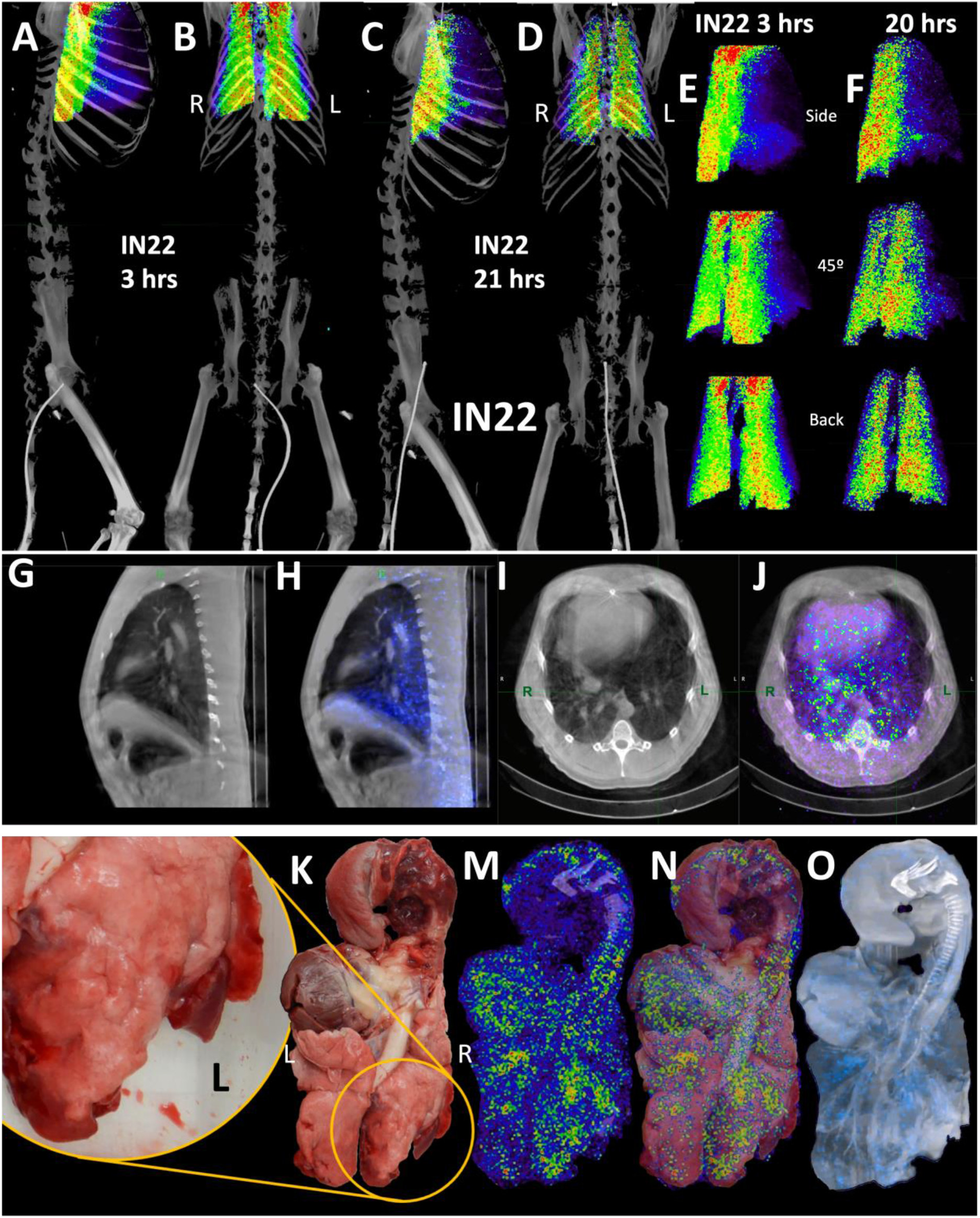
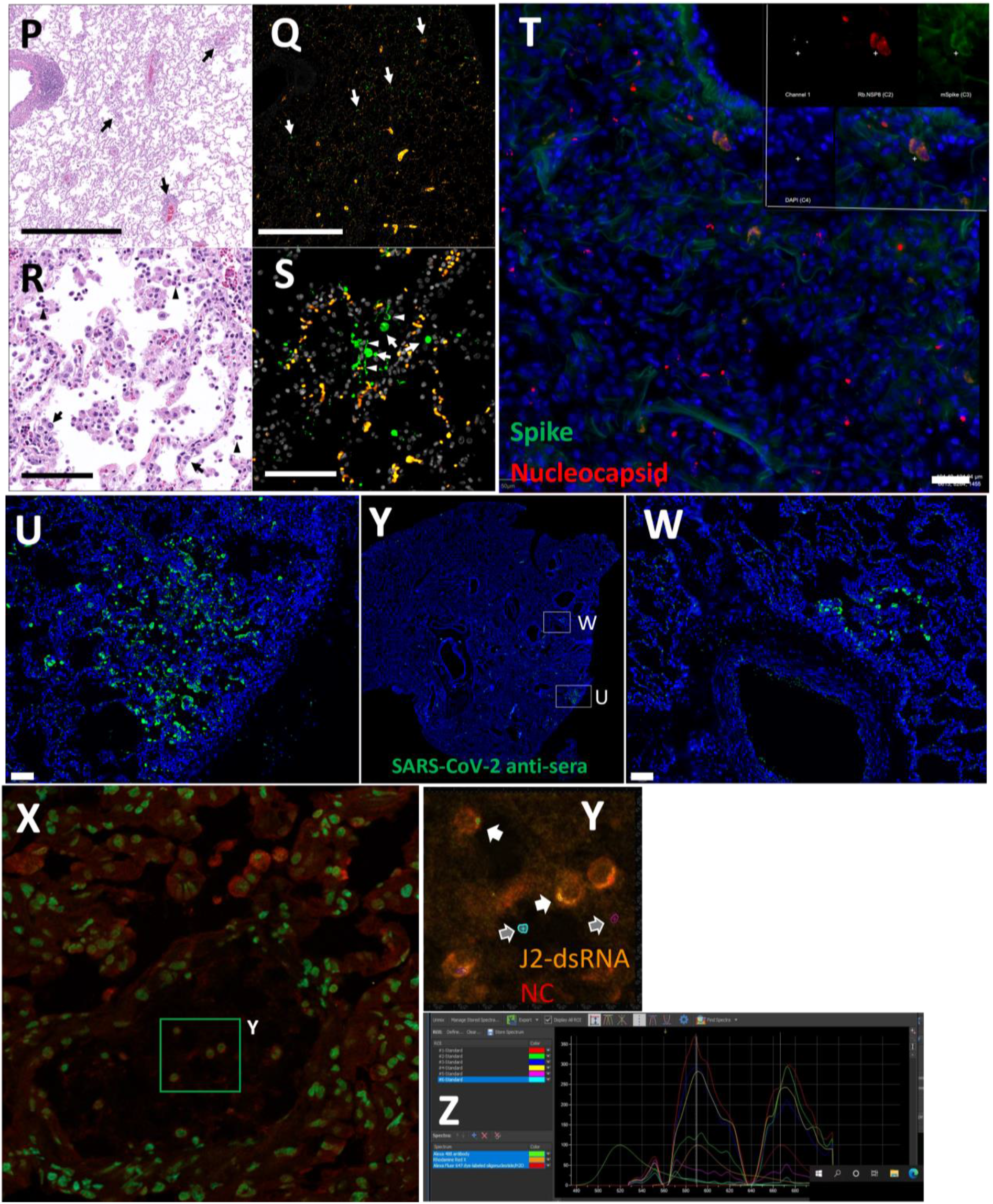
IN22 lung pathology and PET signal. (A-D) Lung PET volumes were isolated and overlaid on whole body CT scans. (A and B) Show lung volumes from 3-hour scan, side (A) and front (B) views shown. (C and D) Show lung volumes from 21-hour scan, side (C) and front (D) views shown. (E and F) Isolated lung PET volumes for each scan are shown independent of CT. (G) Sagittal z-slice from CT showing lungs. (H) PET signal overlaid on z-slice from G. (I) Transverse z-slice through torso from CT. (J) PET signal overlaid on z-slice from I. (K) Image of respiratory tract after necropsy. (L) Inset showing overt lung pathology in right lower lobe. (M) PET/CT signal from organ scan of respiratory tract. (N) PET/CT signal from M overlaid onto image from K. (O) PET/CT signal with CT contrast increased to observe pathology in lower right lung lobe. (P) H&E image of lung tissue showing areas of expanded alveolar space and inflammatory infiltrate (arrows). (Q) Immunofluorescence image of the same tissue from P. Green is SARS-CoV-2 anti-sera, red is background, and white is nuclear stain. White arrows indicate SARS-CoV-2 positive cells. (R) H&E image of lung tissue showing macrophages and neutrophils (arrowheads) and type II pneumocytes (arrows). (S) Immunofluorescence image showing infected cells of the alveoli (arrows) and lining the alveolar septa (arrowheads). Green is SARS-CoV-2 anti-sera, red is background, white is nuclear stain. All scale bars are 100 µM. (T) Fluorescent microscopy image of IN22 lung tissue. Spike shown in green, nucleocapsid shown in red, background in white, and Hoechst nuclear stain in blue. Scale bar 100 µM. (U, Y, W) Additional fluorescent microscopy images of lung tissue showing foci of infected cells. U and W are shown in low magnification Y. Green is SARS-CoV-2 anti-sera and blue is Hoechst nuclear stain. Scales bar 500 µM. (X-Z) Validation of dual antibody staining utilizing spectral imaging. (X) shows microscopy of J2 antibody with redX secondary and rabbit anti-NC monoclonal and Cy5 secondary. (Y) Shows area within green square in X. White arrows point to regions of interest that are cell associated. Grey arrows indicate control regions of spectral evaluation. The areas evaluated by spectral imaging in (Y) are color coded and match with the spectra shown in (Z).

Histopathology of this region revealed marked pulmonary interstitial inflammation and type II pneumocyte hyperplasia (Fig 1D, and 6P-S). The projection of the PET signal over the photo (6N) shows a strong PET signal in multiple lung lobes and located adjacent to the focally extensive areas of consolidation (red and inflamed areas). A similar strong PET signal is observed in several areas of the left lung, with a major PET signal associated with the caudal aspect of the upper lobe of the left lung. An overlay of the unnormalized PET over a CT projection that reveals lung structure and fluid in the lung (Fig 6O) further details the relationship between the areas with evidence of pneumonia and the PET signal. The PET signals tend to be adjacent to opaque regions potentially caused by fluids and the regions of consolidation visible in the photo.

Next, we evaluated paraffin blocks of the right lower lobe by H&E (Fig 6P and R) and immunofluorescent staining for SARS-CoV-2 (Fig 6Q and S). In regions of pneumonia (Fig 6P), the interstitium and alveolar spaces are expanded by inflammatory infiltrate. In these same regions, SARS-CoV-2 infected cells are scattered throughout (Fig 6Q). Higher magnification imaging of the H&E stain (Fig 6R) reveals the inflammatory infiltrate is composed predominately of macrophages with lesser neutrophils (arrowheads). Alveolar septa are frequently lined by type II pneumocytes (arrows). SARS-CoV-2 staining (Fig 6S) reveals infected cells within alveoli (arrows) and lining alveolar septa (arrowheads).

To validate the accuracy of our SARS-CoV-2 staining results, 2-color immunofluorescence staining for spike and nucleocapsid was used to identify SARS-CoV-2 infected cells in the right lung (Fig 6T). Spectral imaging confirmed that the fluorescence signal associated with the double positive cells is consistent with the specific fluorophores utilized for antigen visualization. Additional fluorescent staining shows focal nature of infected cells in a large piece of lung tissue (Fig 6U-W). The 2-color imaging is a powerful method to validate the identification of SARS-CoV-2 infected cells. Spectral imaging of left lung tissue stained for NC and dsRNA was used as an alternate approach to identify infected cells (Fig is shown in panels 6X-Z). The spectrum shown in Fig 6Z confirms that the cells identified in panel 6Y are specifically double stained with the antibodies to dsRNA and SARS-CoVCoV-2 NC.

To evaluate the probe signal in the MGT after Delta infection, we compared the 3-hour and 21-hour PET/CT signal in the IN22 MGT (Fig 7A-D); the isolated PET signal of the MGT of both scans is shown below (Fig 7E-H). A signal is apparent in the base of the testes that becomes more diffuse at the 21-hour timepoint. There is also a signal apparent just above both testes in the 3-hour scan that increases at 21-hours and is most prominent above the right teste (Fig 7A-H, indicated by white asterisk). Examination of both scans indicates that the probe signal was initially associated with the vasculature, but it persisted and accumulated adjacent to the top of testes in the 21-hour scan. This PET signal is associated with the vasculature of the spermatic cord as is visible in the CT projections and the PET overlays in the 21-hour scan (Fig 7I-L) and after the necropsy (Fig 7M-P). The post necropsy signal matches the *in vivo* scan with the left testicle having a persistent signal throughout the spermatic cord and a more localized signal on the top of the right teste. To confirm infection in these tissues, the tissue block with the greatest PET signal (L Test 1, Fig 5I) was sectioned for genomic and microscopic analysis. Bulk RNA was isolated from 2 tissue sections and qPCR analysis of the SARS-CoV-2 N gene revealed the presence of viral RNA (2.03 Copies N1/ul). Immunofluorescence analysis of the PCR positive tissue derived from the same block revealed the presence of sparse cells double positive for spike and nucleocapsid primarily found in the interstitial space (Fig 7Q and R). Staining of penile tissue from an uninfected macaque (Fig 7S) reveals robust ACE2 expression in the venous spaces of the corpus cavernosum. Multiple SARS-CoV-2 infected cells in the penis were revealed by double staining for nucleocapsid and dsRNA (J2) (Fig 7T-W).

**Figure 7.**
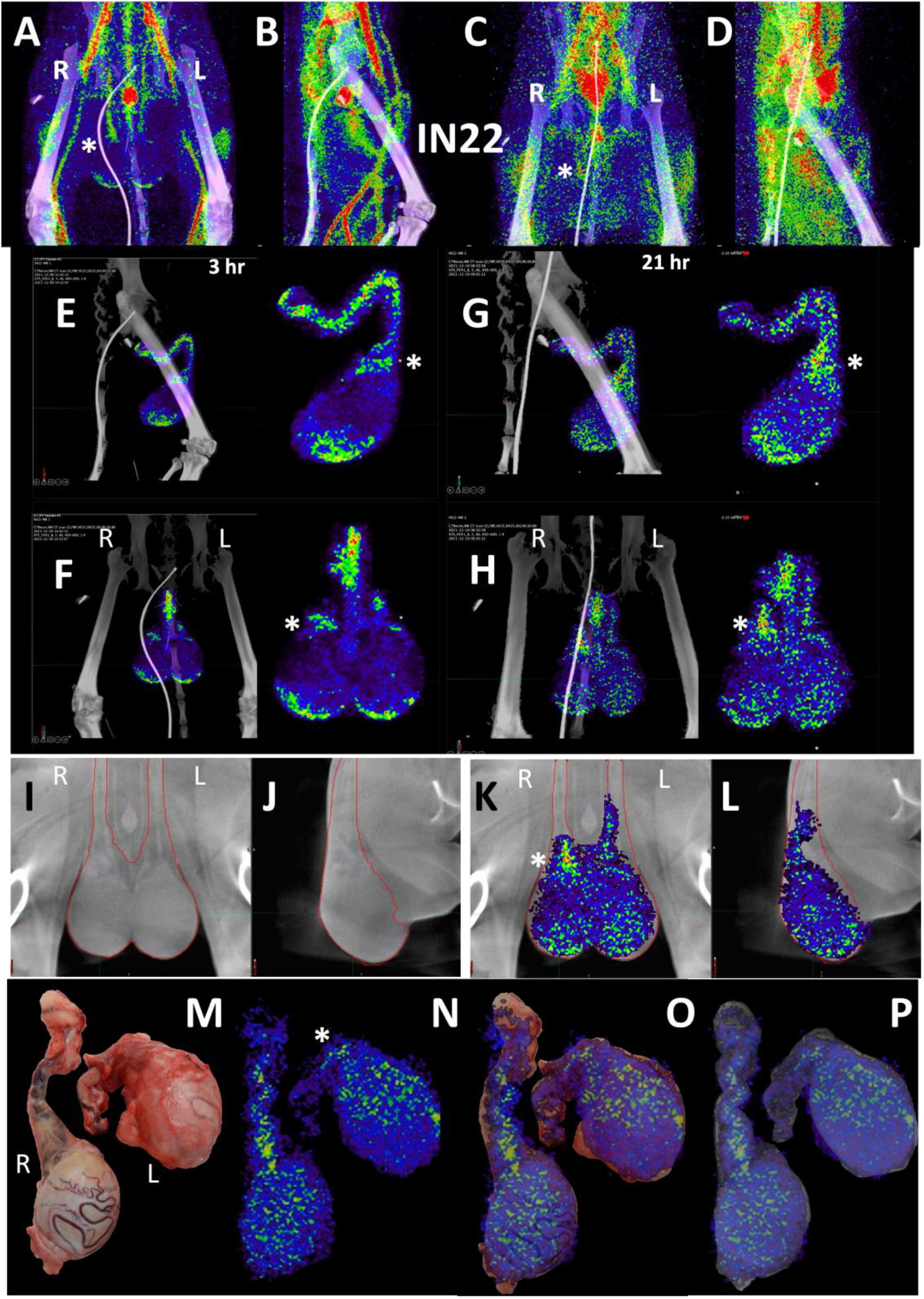
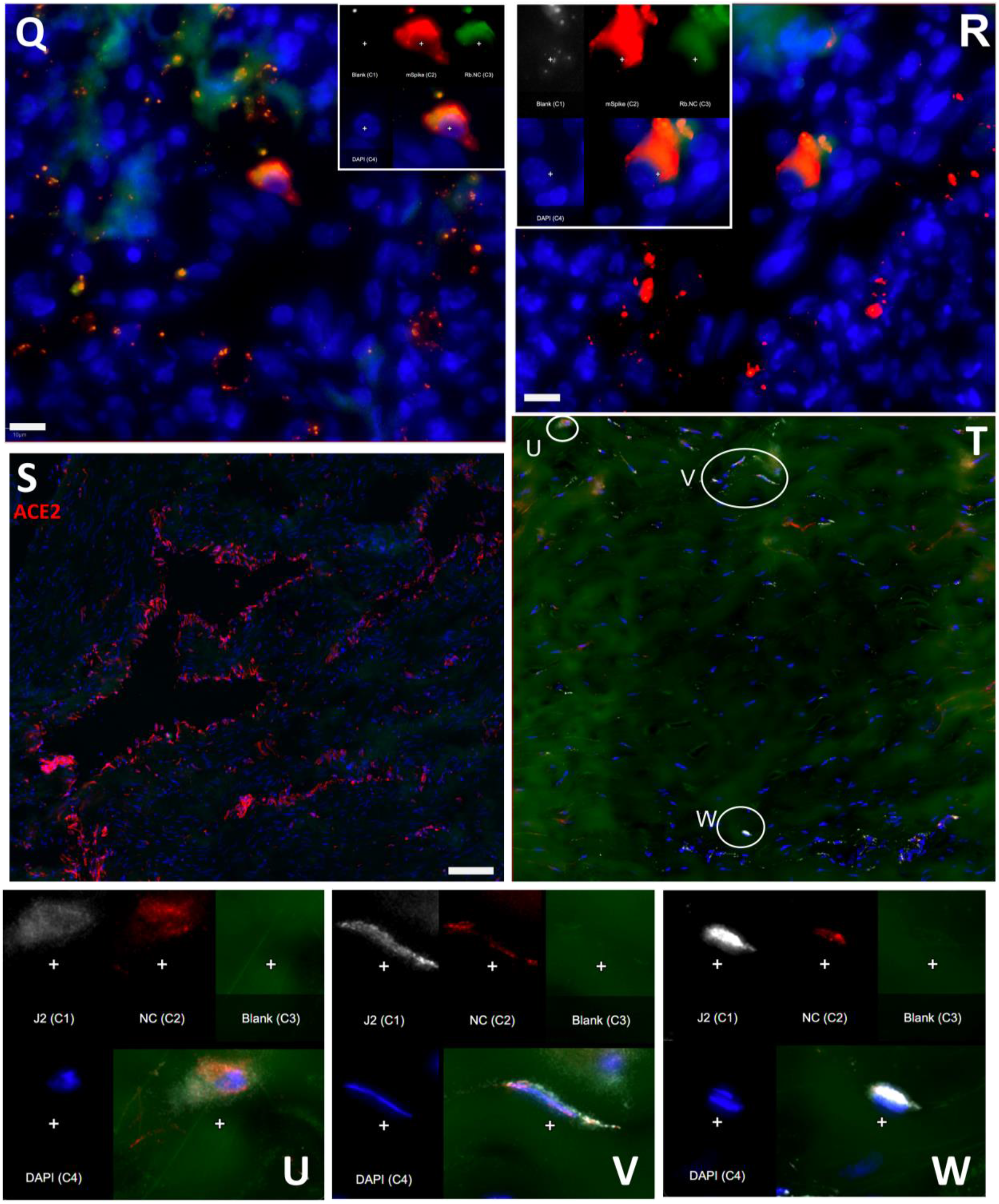
Male genital tract signal of IN22. (A and B) PET/CT images highlighting the lower abdomen of IN22 obtained 3-hours after probe administration. Front view (A) and side (B) both shown. (C and D) PET/CT images highlighting the lower abdomen of IN22 obtained 21-hours after probe administration. Front view (C) and side (D) both shown. (E and F) Isolated 3D volume of MGT from 3-hour scan overlaid with whole-body CT. Side (E) and front (F) views shown. (G and H) Isolated 3D volume of MGT from 21-hour scan overlaid with whole-body CT. Side (G) and front (H) views shown. (I and J) Whole-body CT images of lower abdomen, red contours outline the testicles and spermatic cords. Front (I) and side (J) views shown. (K and L) 3D volume of MGT overlaid onto CT images from previous panels. (M) Image of testicles after necropsy. (N) PET signal from organ scan of testicles. (O) Overlay of PET signal onto image from panel M. (P) Overlay of PET signal onto CT signal from same scan. White asterisks mark location of pampiniform plexus in all previous panels. (Q and R) Fluorescence microscopy of SARS-CoV-2 infected cells in testicular tissue from IN22. Red is SARS-CoV-2 spike, green is SARS-CoV-2 nucleocapsid, white is background, and blue is Hoechst nuclear stain. Insets show channels independently, larger image is all channels merged. Scale bars 10 µM (S) Fluorescent microscopy image of corpus cavernosum tissue from an uninfected animal showing ACE2 staining in red, background in green, and Hoechst nuclear staining in blue. Scale bar 100 µM (T) Fluorescence microscopy of SARS-CoV-2 infected cells in penile tissue from IN22. White is dsRNA antibody J2, red is SARS-CoV-2 nucleocapsid, green is background, and blue is Hoechst nuclear stain. (U, V, and W) Show individual cells highlighted in T.

### Longitudinal analysis of SARS-CoV-2 Delta variant distribution 1- and 2-weeks post-infection

For the final pilot study, we performed two longitudinal PET/CT scans, at 1- and 2-weeks post challenge with the Delta variant, on a single animal (JF82). The goal of this study was to determine if the ^64^Cu-F(ab’)2 probe could provide novel insights into the spatiotemporal dynamics of SARS-CoV-2 infection with sequential PET/CT scans. A front and side view of the sequential PET/CT scans at 1-week (Fig 8A-B) and 2-weeks (Fig 8C-D) reveals dynamic changes in various organ systems. For example, there is a decrease in the signal of the lungs between week one and two while MGT signal increases. These changes are illustrated in the front (Fig 8E) and side (Fig 8F) view overlay of the 1-week (shown in red) and 2-week (shown in blue) timepoints. Areas with blue signal indicate increased signal in week 2 relative to week 1 and areas with red signal indicate where the week 1 scan had greater signal. Areas with white signal indicate where there is high PET signal in both scans. The post necropsy organ scan (Fig 8G-J, S7-video) and the tissue scan (Fig 8K) similarly show probe signal associated with both the lungs and MGT.

**Figure 8.**
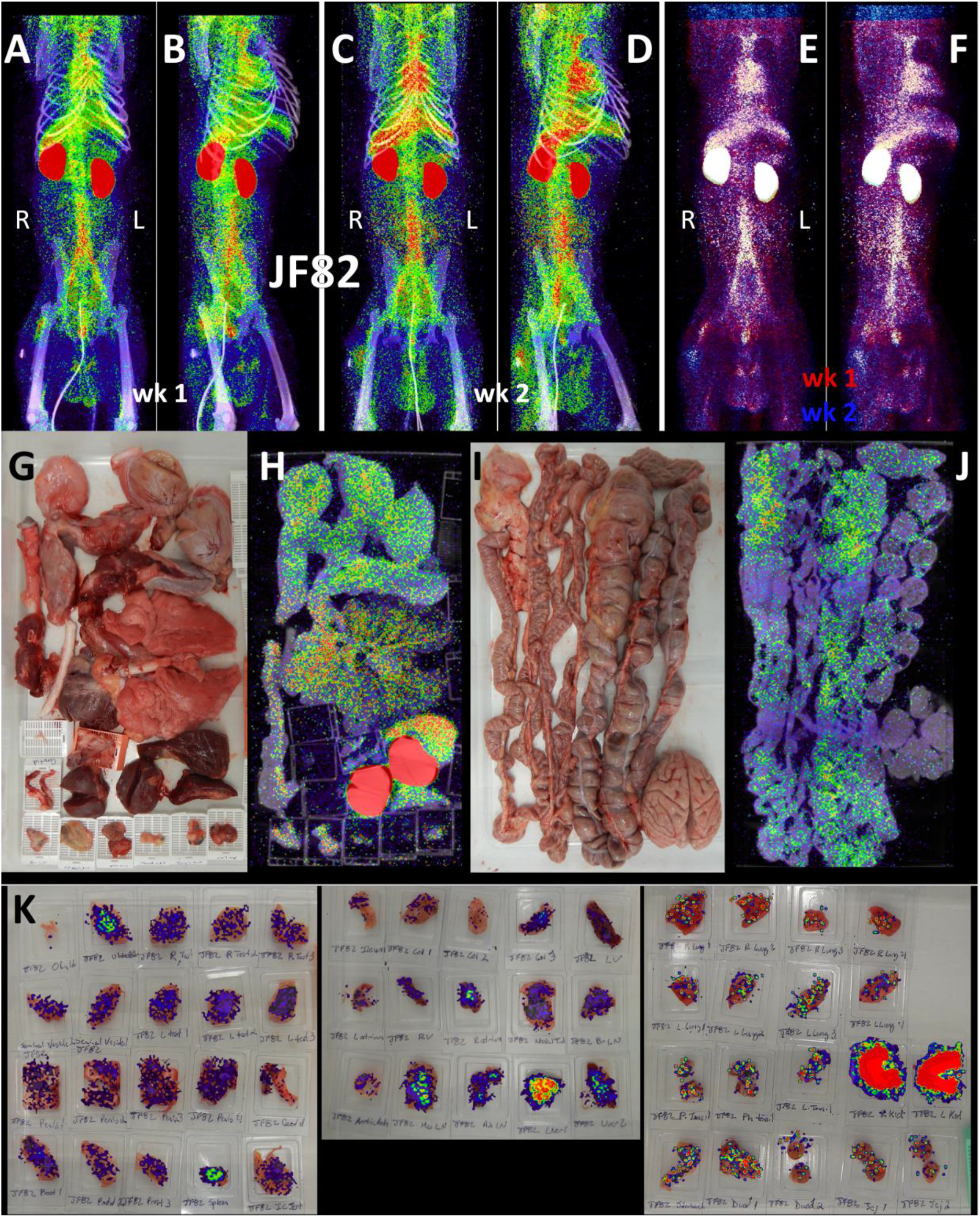
JF82 PET/CT images from week and week 2 necropsy scans. (A and B) Whole-body PET/CT scan of JF82 from 1-week post-infection. Front view (A) and side (B) both shown. (C and D) Whole-body PET/CT scans of JF82 2-weeks post infection. Front view (C) and side (D) both shown. Right and left labeled in front views (A and C). (E and F) Overlay of the week 1 scan (shown in red) and the week 2 scan (shown in blue). Front (E) and side (F) views both shown. (G) Organ tray post necropsy and (H) PET/CT image. (I) Second organ tray post necropsy and (J) PET/CT image. (K) Overlay of PET signal onto photograph of tissue cassettes.

To better compare the probe signal in the lungs of JF82 at the 1-week (Fig 9A-B) and 2-week (Fig 9C-D) timepoints, the 3D reconstruction of the JF82 lungs was isolated from the PET datasets and projected over the CT reconstruction of the skeleton (Fig 9A-D). Both the 1- and 2-week lung signals are apparent and localized with a level of signal comparable to the previous animal (IN22) infected with the Delta variant. An evaluation of the data set revealed a PET signal overlying a region of opacity in the lower lobe of the left lung as designated with the asterisks in several of the panels (Fig 9A-B, 9G-L). The lung rotation series shown for week 1 (Fig 9E) and week 2 (Fig 9F) reveal a major signal associated with the dorsal side of the left lung at both timepoints and less signal associated with the right lung in the week 1 scan. Evaluation of signal from coronal sections of the week 1 PET/CT overlay (Fig 9G) and CT alone (Fig 9I) reveals an overlap of the probe signal with an opaque region consistent with focal pneumonia. It is notable that both the PET and CT signal associated with this spot in the left lung are gone in the week 2 scan (Fig 9H and J). This is consistent with reports that the lung pathology observed in the rhesus macaque model is most apparent after 1 week of infection and can wane by week 2^25^. To better illustrate the change between week 1 and week 2, the week 1 scan in red and the week 2 scan in blue were overlaid with the week 1 CT signal (Fig 9K and L). Microscopic analysis revealed pulmonary infiltrates were still present in the alveolar space at necropsy (Fig 1E). However, no infected cells were detected with immunofluorescence using an anti-SARS-CoV-2 antibody in FFPE tissue. These findings are suggestive of a resolving infection which is supported by the histopathology (Fig 1E) and viral RNA levels (Fig 1F and G).

**Figure 9.**
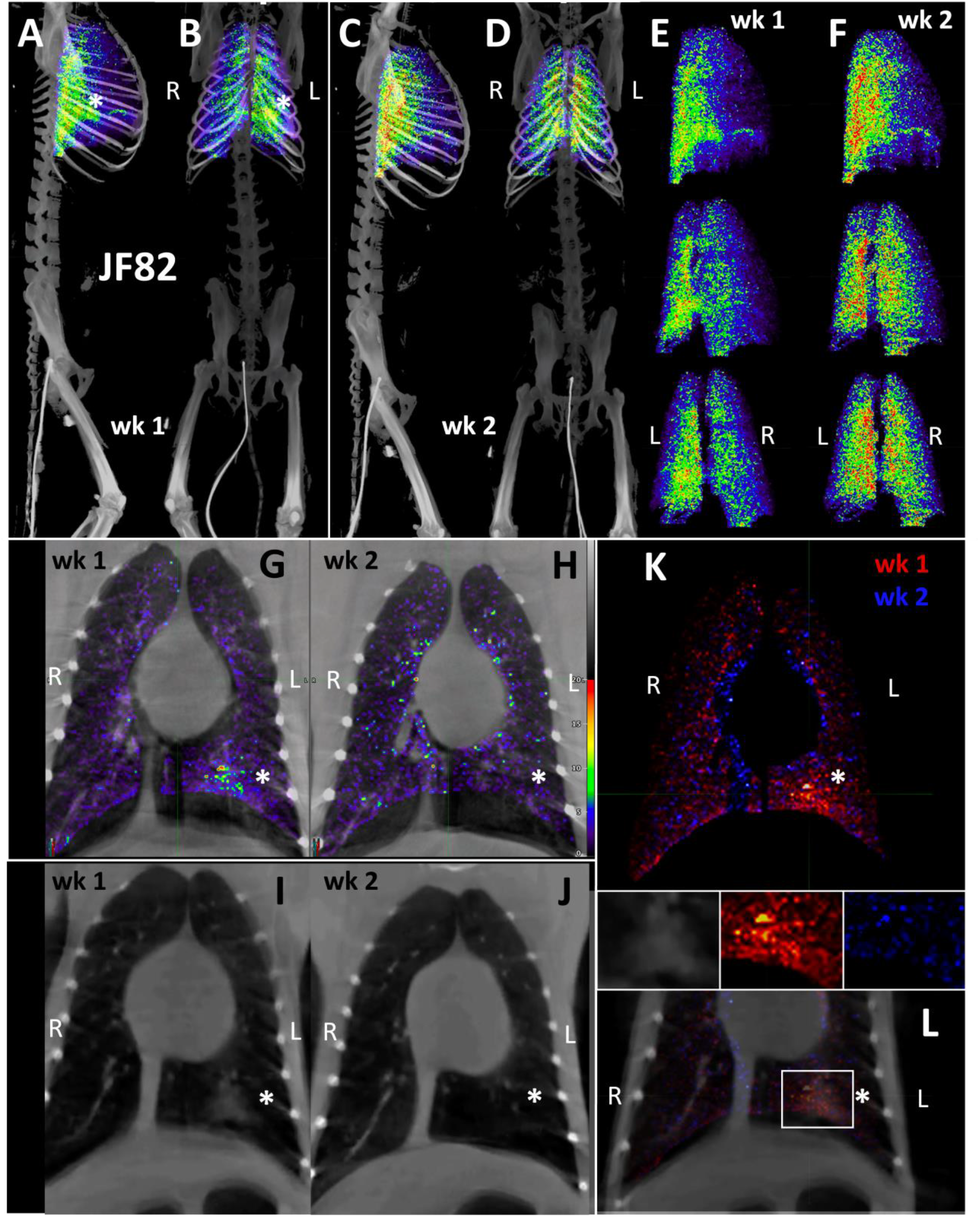
JF82 lung pathology and PET signal from two timepoints. (A-D) Lung PET volumes were isolated and overlaid on whole body CT scans. (A and B) Show lung volumes from 1-wee scan, side (A) and front (B) views shown. White asterisk indicates location of lung pathology highlighted below. (C and D) Show lung volumes from 2-week scan, side (C) and front (D) views shown. (E and F) Isolated lung PET volumes for each scan are shown independent of CT. (G and H) Overlay of single z-image of PET signal onto single z-image of CT in the lungs at week 1 (G) and week 2 (H). (I and J) Single z-image of CT used in G and H shown independent of PET signal for week 1 (I) and week 2 (J). (K) Overlay of week 1 (shown in red) and week 2 (shown in blue) PET signal. (L) Overlay from K shown with CT image to localize PET signal. Insets above show, CT, week 1, and week 2 signal from left to right. White asterisk indicates location of left lung pathology in all panels.

We next evaluated the signal associated with the MGT of JF82 as illustrated in Fig 10, which presents the front and near side view (∼45°) of the abdominal area of the week 1 (Fig 10A and D), week 2 (10B and E), and overlay (10C and F). The overlay (Fig 10C and F) reveals the dynamics of the probe signal in the MGT of JF82 in the first 2 weeks of SARS-CoV-2 infection. The white signal in the overlay reveals that the probe signal is maintained in the prostate, the vasculature at the base of the spermatic cord, and the base of the testes. To gain additional insights into the MGT associated signal at the 2 time points, we isolated the MGT volumes and 3D projected the signal (Fig 10G-J). In the week 1 scan (Fig 10G-H), a signal associated with the root of the penis is also apparent in addition to the signal associated with the vasculature at the base of the spermatic cord and the base of the testes. In the week 2 scan, the signal becomes more diffuse, spreading throughout the penis and testes, and extending into the spermatic cord, especially into the right spermatic cord. To better visualize the signals associated with the different tissues, we isolated the volumes containing the penile signal for the week 2 scan (Fig 10K-M). The signal distribution throughout the penis at week 2 is readily apparent and distinct from the signal associated with the spermatic cord. It is notable that the probe signal associated with the MGT becomes better distributed and more pronounced in the week 2 scan, consistent with a spreading infection in the MGT between week 1 and week 2. In contrast, a focus of infection in the right lung (Fig 9) of the same animal is observed in the week 1 scan and resolved in the week 2 scan.

**Figure 10.**
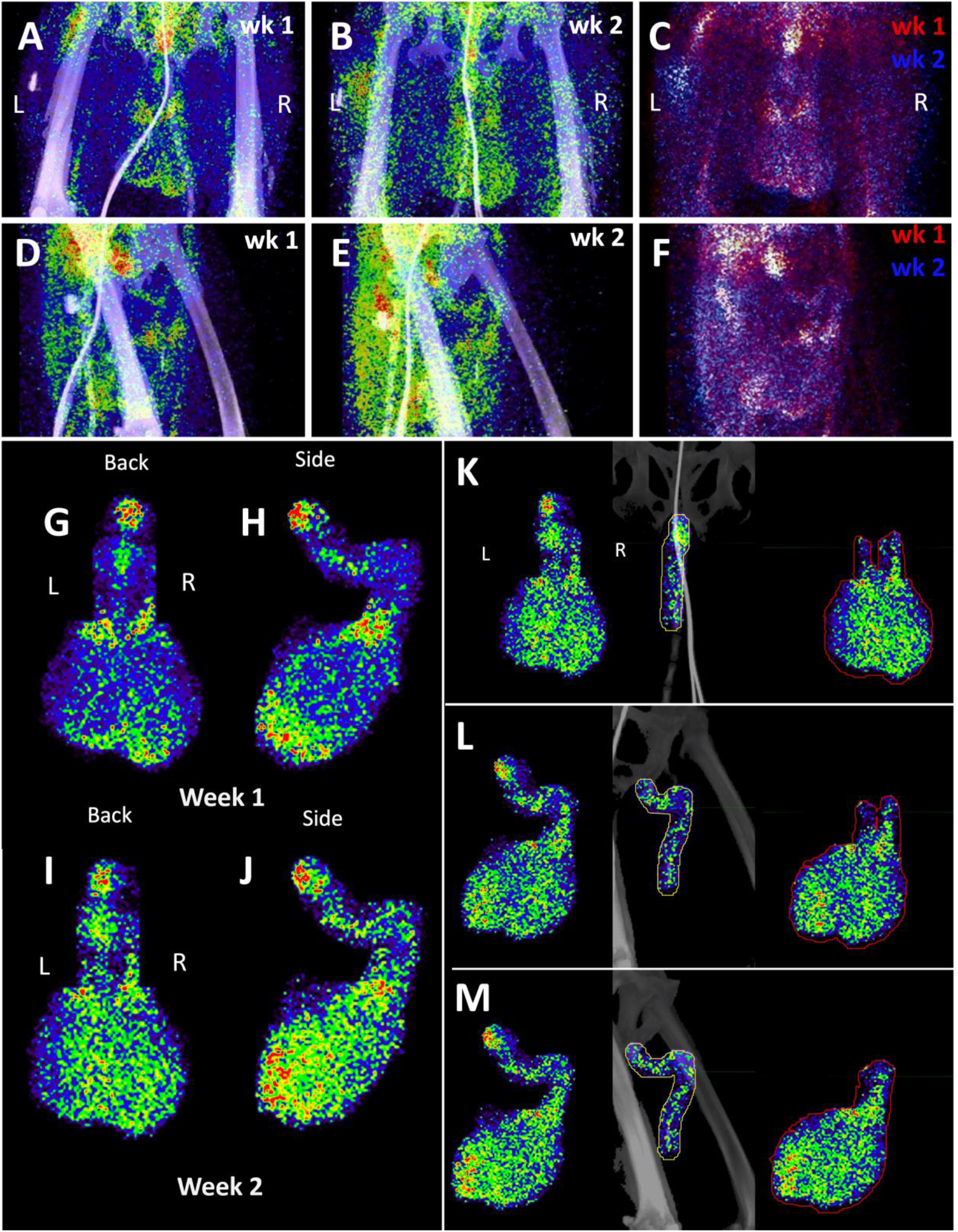
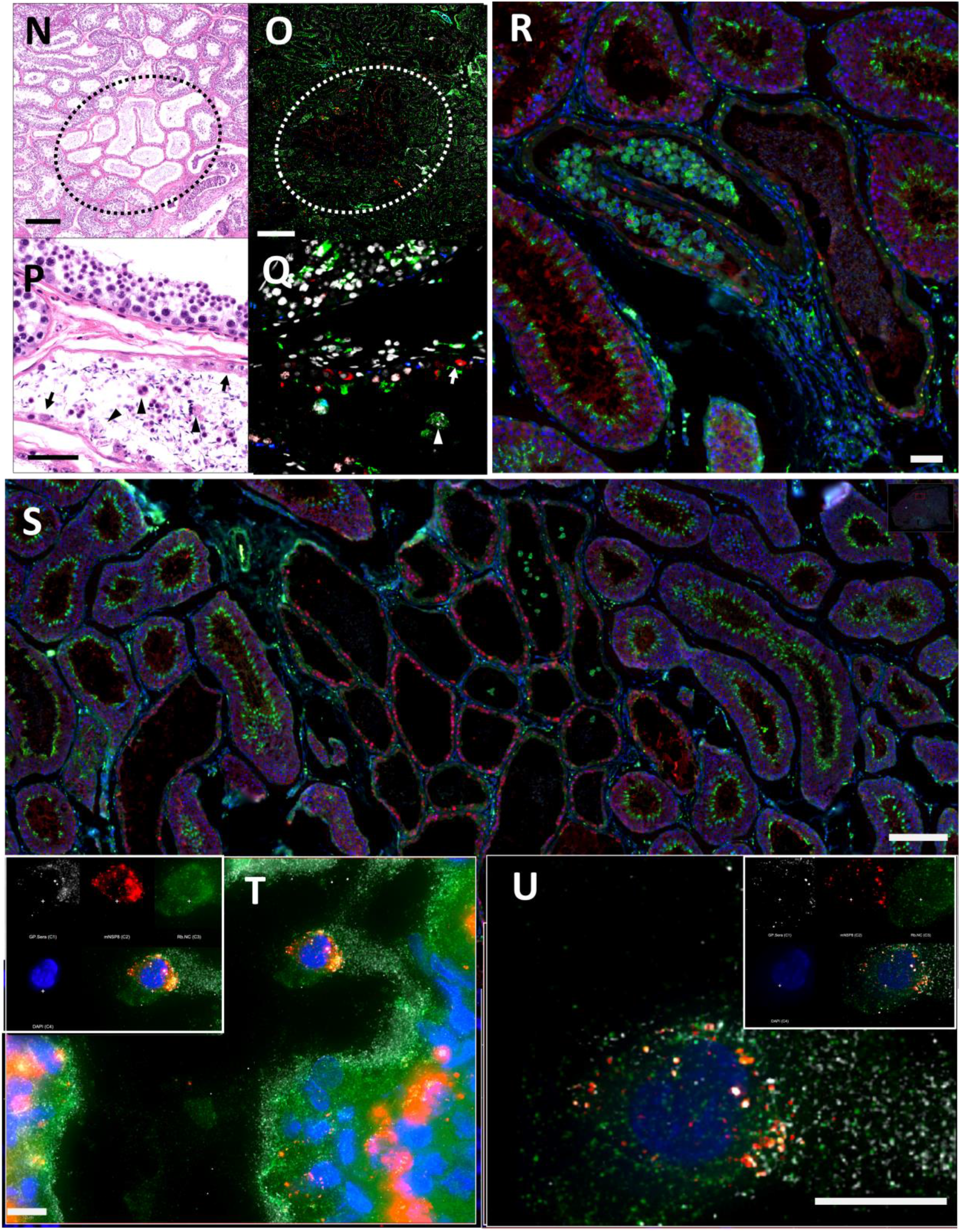
Male genital tract signal in JF82. (A and B) Front view PET/CT images highlighting the lower abdomen of JF82 at 1-week (A) and 2-weeks (B). (C) Overlay of week 1 (shown in red) and week 2 (shown in blue) PET signal. (D and E) Side view of same images shown in A and B. (F) side view of overlay shown in C. (G and H) Isolated MGT volume from week 1 scan. Back (G) and side (H) views shown. (I and J) Isolated MGT volume from week 2 scan. Back (I) and side (J) views shown. (K, L, and M) Isolated MGT volume from week 2 with isolated penile and testicular signal. Back (K), rotated 45° (L), and side (M) views shown. (N) H&E stain of JF82 testicular tissue. Degenerate seminiferous tubules highlighted in black oval. Scale bar 500 µM. (O) Fluorescent microscopy shows a similar area of degenerate tubules. Green is CD206, red caspase 3, and white nuclear stain. Scale bar 500 µM. (P) Higher magnification image of degenerate tubules. Intraluminal macrophages (arrowheads) and Sertoli cells (arrows) are shown inside tubules. Scale bar 50 µM (Q) Higher magnification of image in O shows Sertoli cells (arrow) staining with caspase 3 and macrophages (arrowhead) staining with CD206. Scale bar 50 µM. (R) Fluorescent microscopy image showing degenerate tubule full of intraluminal macrophages. Red is caspase 3, green is CD206, and blue is nuclear stain. Scale bar 50 µM. (S) Lower magnification microscopy image of degenerate and healthy tubules. Red is caspase 3, green is CD206, and blue is nuclear stain. Scale bar 200 µM. (T and U) Fluorescent microscopy images of infected cells in testicular tissue of JF82. White is SARS-CoV-2 anti-sera, red is NSP8, green is nucleocapsid, and blue is Hoechst nuclear stain. Insets show individual channels, larger image is merge. Scale bars 10 µM.

Histopathology of testicular tissue from JF82 revealed multifocal regions of degenerate seminiferous tubules characterized by a complete loss of germ cells and spermatids (Fig 10N). These regions also have evidence of edema as revealed by increased spaces between individual seminiferous tubules. Degenerate seminiferous tubules occasionally contained macrophages with phagocytosed spermatids, and the adjacent interstitium was infiltrated by low numbers of lymphocytes and plasma cells. To further characterize the degenerative changes noted on H&E, immunofluorescence for CD206 - a mannose receptor present on monocytes, macrophages^27^, and mature spermatids^28–30^ - and caspase 3 - a cellular marker of apoptosis - was performed. Degenerate seminiferous tubules were readily identified by the marked decrease in CD206 expression (due to loss of mature spermatids) and increased expression of caspase 3 compared to adjacent, nondegenerate, tubules (Fig 10O, Q-S). Evidence of intra-tubule macrophages was readily apparent (Fig. 10R, S). SARS-CoV-2 infected cells can be identified in the JF82 testes with triple staining for NSP8, nucleocapsid, and SARS-CoV-2 anti-sera (Fig 10T and U).

### Comparison of PET signal between animals

A comparison of the PET/CT scans of 3 SARS-CoV-2 infected rhesus macaques is revealing in terms of the dynamics of infection and the utility of this technique to study COVID-19. In Fig 11A-D we present a comparative rotation series of the PET signal withing the lung volumes of all scans. Although there are differences in the lung signal of each scan, the Delta variant utilized for infection of IN22 (Fig 11B) and JF82 (Fig 11C-D) is associated with increased pathology^26^ (Fig 1C-E, Fig 5, and Fig 8) and increased PET signal (Fig 11B-D) without a clear difference in viral load in nasal swab or saliva at 1 week after challenge (Fig 1F and G) compared to LP14. This is consistent to similar viral loads between WA1 and the Delta variant in a recent report^26^.

**Figure 11.**
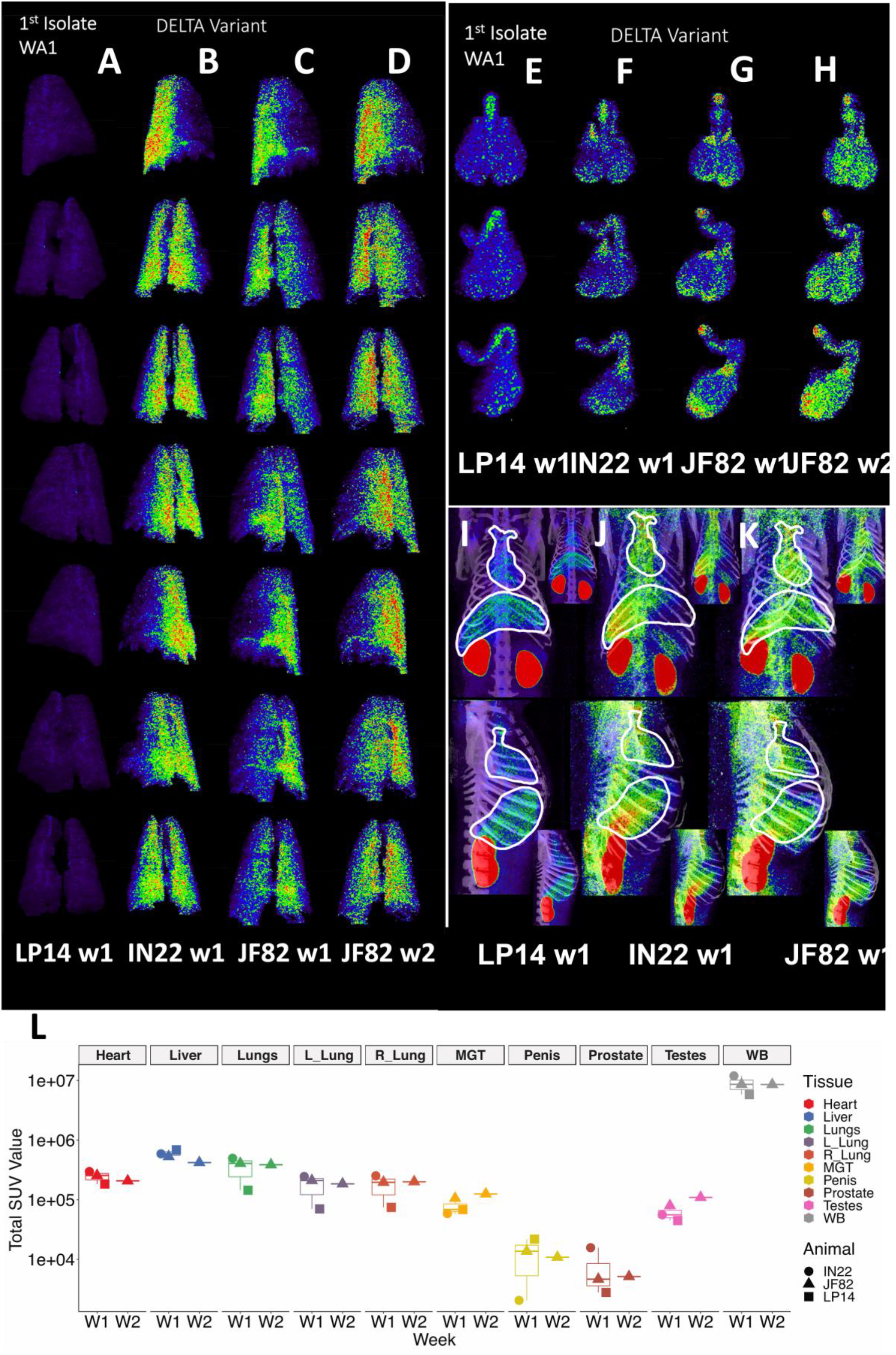
Comparison of SUVs across animals and timepoints. (A-D) Complete rotation series of lung PET volumes for LP14 (A), IN22 (B), JF82 week 1 (C), and JF82 week 2 (D). (E-H) Front, rotated 45°, and side views of MGT PET volumes for LP14 (E), IN22 (F), JF82 week 1 (G), and JF82 week 2 (H). (I-K) Front and side views of whole-body scans, white lines indicate volumes taken for heart and lungs for LP14 (I), IN22 (J), and JF82 week 1 (K). Insets show each image without white outlines. (L) Total SUVs for whole-body scans and each individual volume isolated displayed in graph. Animals are indicated by icon shape and volumes by color.

The probe signal associated with the MGT was not anticipated, but apparent in all 3 animals. This reveals that infection of MGT is consistently seen in the rhesus macaque model of SARS-CoV2 IN/IT challenge. A comparison of the isolated MGT PET signal from the four whole-body PET scans of 3 SARS-CoV-2 infected rhesus macaques (Fig 11E-H) presented as a rotation series further reveals the dynamics of SARS-CoV-2 after infection. The difference in MGT signal between the animal infected with WA1 (LP14) and the animals infected with the Delta variant (IN22 and JF82) at week 1 is much less pronounced than that seen in the lungs. In all week 1 MGT scans, the signal is asymmetrically distributed with diffuse signal throughout the testicles, an increased signal at the base and top of the testes, and a signal associated with the root of the penis (Fig 11E-G). The MGT PET signal is visibly increased in the week 2 scan relative to the week one scan (Fig 11G, H) for JF82 indicating further spread of infection into the MGT at that time. Another obvious difference between the week 1 whole-body PET scans is a variable signal associated with the heart and liver as shown in Fig 11I-K. LP14 had a diffuse PET signal throughout the liver (Fig 11I). In contrast, the IN22 PET signal (Fig 11J) was primarily localized with the right side of the liver and in JF82 the PET signal (Fig 11K) was localized to the base of the liver. Additionally, the extent of labeling of the heart is variable with JF82 and IN22 having a greater signal than LP14 (Fig 11L).

To take advantage of the quantitative aspects of PET detection, we isolated the total standard uptake value (SUV) of the PET signal associated with the CT defined volumes as plotted in Fig 11L. The 3 animals scanned 1 week (W1) post SARS-CoV-2 challenge are presented together for the whole-body (WB) scan signal and all evaluated tissues. The single week 2 (W2) PET signal of JF82 is presented for comparison. The WB values for all scans are clustered revealing the reproducible nature of evaluation of the PET signal. An increase in the total SUV between week 1 and week 2 for the MGT and testes volumes is consistent with the increase of signal suggested by visual inspection of the isolated tissues (Fig 10, Fig 11G and H).

Another relevant tissue with PET signal observed in all 3 animals is the prostate. Probe labeling of the prostate first became apparent in the early PET scan (3 hr. after injection) of IN22 where it was among the strongest, non-kidney associated signals (Fig 5A and B). Therefore, we revaluated the PET/CT data sets for the 3 animals. We were able to detect a signal associated with the prostate in all animals (marked by white asterisks, Fig 12A-F). An inset showing a sagittal slice of the PET/CT shows the localization of the prostate within the small white circle for each animal. PET signal was also isolated within the penile volume of the whole-body scan for IN22 1-week post infection (Fig 12G-I) and 2-weeks post infection for JF82 (Fig 12J-L). LP14 is excluded from this analysis because the penile tissue was not collected. The same general distribution of PET signal is seen in the organ scan of IN22 (Fig 12M) and JF82 (Fig 12N). The localization of the PET signal is further defined in the tissue scan of the cryomolds shown with (Fig 12O and Q) and without (Fig 12P and R) PET signal for each animal. There is PET signal associated with the penile tissue with a stronger signal associated with the penile root of IN22 and the glans of JF82. This PET/CT guided necropsy of the MGT reveals a persistent signal associated with the penile tissue of these 2 animals. All 3 animals had an apparent PET signal associated with the testes at the 1-week timepoint and that signal increased in the testes in the week 2 scan (Fig 12Q-T, quantified in Fig 11O). The histopathologic analyses of the testes from all three animals are shown in Fig 12S-U. Testicular degeneration was noted in seminiferous tubules of JF82 (Fig. 10N). Panels demonstrate normal spermatogenesis or lack thereof in JF82 (Fig. 12N, O, R, S).

**Figure 12.**
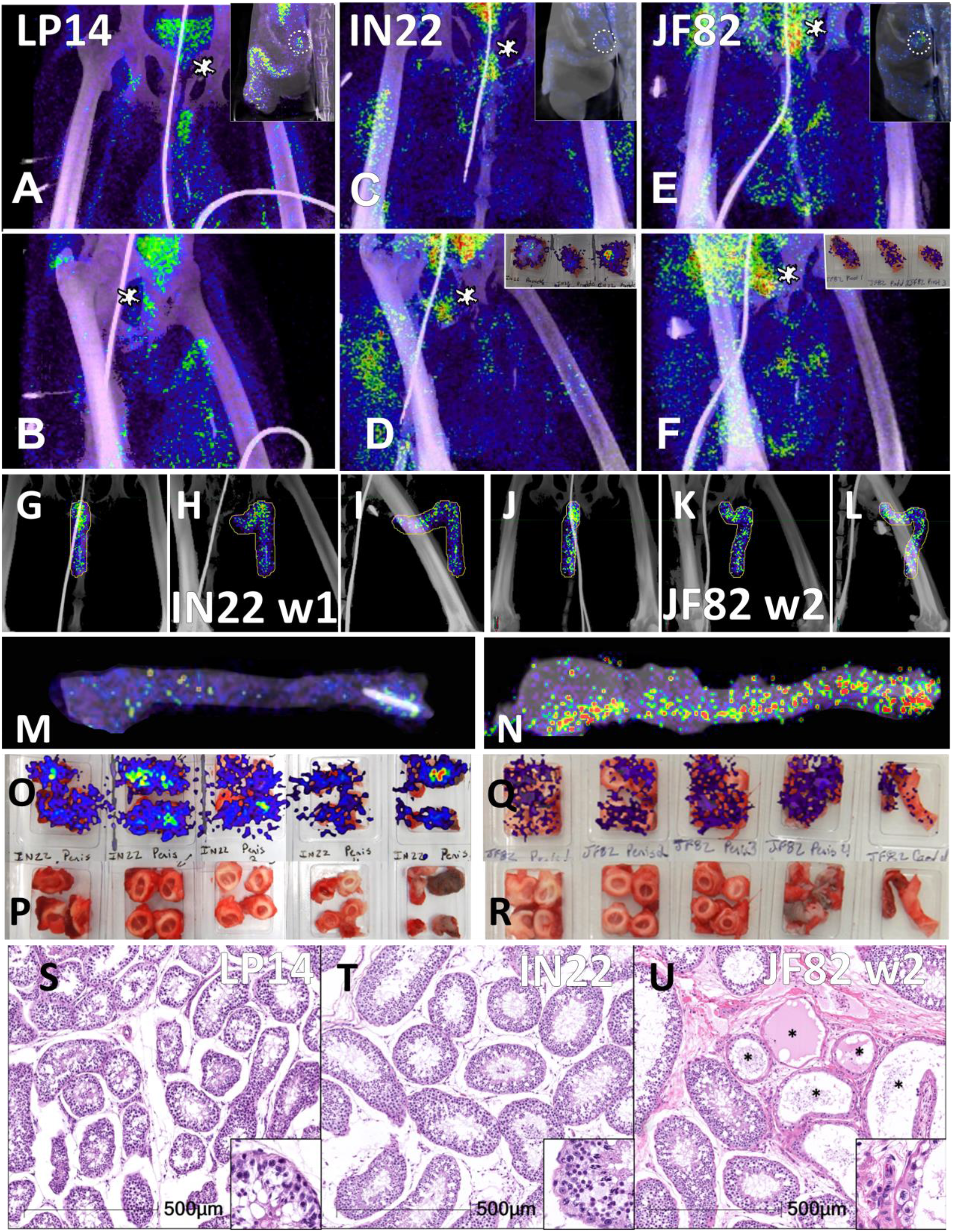
Comparison of prostate and penile signal between animals. (A-F) PET/CT signal in lower abdomen for each animal at 1-week post-infection. LP14 front (A) and rotated 45° (B), IN22 front (C) and rotated 45° (D), JF82 front (E) and rotated 45° (F). Asterisks mark location of prostate. Insets show sagittal z-slice of each animal highlighting prostate signal (white circle). (G-I) PET/CT volume of penis for IN22. Front (G), rotated 45° (H), and side (I) views shown. (J-L) PET/CT volume of penis for JF82 at 2-weeks post-infection. Front (J), rotated 45° (K), and side (L) views shown. (M) PET/CT signal of IN22 penis after necropsy. (N) PET/CT signal of JF82 penis after necropsy. (O) PET/CT signal overlaid onto an image of tissue cassettes (P) containing penile tissue of IN22. (Q) PET/CT signal overlaid onto an image of tissue cassettes (R) containing penile tissue of JF82. (S-U) H&E images of testicular tissue from each animal. LP14 (S) and IN22 (T) shows normal spermatogenesis and tissue architecture. IN22 (U) shows degenerate tubules (asterisks) interspersed among healthy tubules. Scale bars all 500 µM.

In addition, all animals had a PET signal located at the top of each testicle, where the spermatic cord connects with testicle. A dissection of the rhesus macaque testicular anatomy is shown in Fig 13A-C. The spermatic cord contains the vasculature supplying blood to the testicles, the vas deferens which transports mature sperm produced in the testes, and the cremaster muscle (Fig 13A). The position in natural context of the macaque penis, testes, and spermatic cord are shown in Fig 13B. A magnified view of the vasculature of the pampiniform plexus is shown in Fig 13C. As shown in the IN22 MGT PET/CT series (Fig 13D-G), there is a major signal associated with the top of the testicles, especially the right testicle. A further examination of this signal within the PET/CT data set demonstrates it is located just above the testicle in the yellow volume (Fig 13D-G). This yellow volume is overlapping with a vasculature structure consistent with the pampiniform plexus visualized by CT (Fig 10G and B). The signal associated with the pampiniform plexus and spermatic cord is seen in the 1-week scan of all animals in front (Fig 13H, J, and L) and rotate 45° view (Fig 13I, K, and M) outside of the ovals indicating the tetes. The signal associated with the pampiniform plexus and spermatic cord for LP14 can be seen outside of the white ovals (Fig 13L-Q).

**Figure 13.**
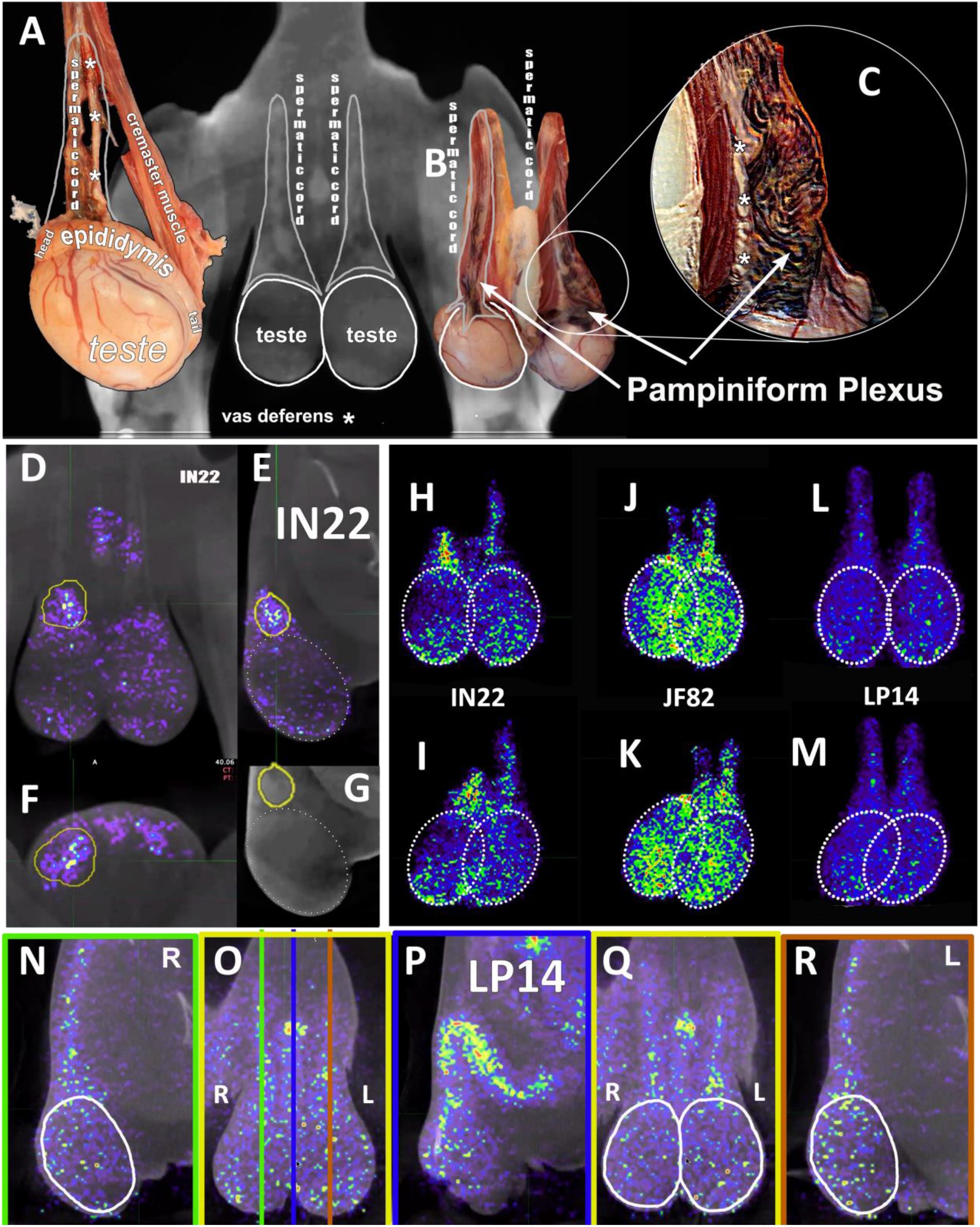
PET signal associated with the pampiniform plexus. (A) Labeled dissection showing anatomical structure of a macaque testis. (B) CT image of testes and associated image showing the matching anatomical features of the spermatic cord. (C) Inset highlighting the location and appearance of the pampiniform plexus. White asterisks mark the vas deferens in all images. (D-F) Single z PET/CT images of IN22 highlighting the PET signal associated with the pampiniform plexus (yellow volume) from the frontal (D), sagittal (E), and transverse (F) plane. (G) CT image used in E to highlight signal associated with pampiniform plexus (yellow volume). (H-L) Isolated 3D volumes of MGT PET signal shown from the front for IN22 (H), JF82 (J), and LP14 (L). White circles outline testes. (I, K, M) Volumes from H, J, and L rotated 45°. (N) Sagittal z-slice of PET/CT of LP14 showing right testis. (O) Frontal z-slice of PET/CT, colored lines correspond to sagittal slices shown in N, P, and R. (P) Sagittal z-slice of PET/CT showing penile tissue. (Q) Frontal z-slice highlighting the testicular tissue in white ovals. (R) Sagittal z-slice of PET/CT showing left testis of LP14. White ovals highlight signal associated with testes and not pampiniform plexus.

### Principal component analysis of PET SUV signals

The PET signal includes several parameters in each anatomical area, each of which provide different insights into the distribution of the probe within the tissue. For example, the total SUV (Fig 11L), provides insights into the overall signal in each tissue/animal. However, the total SUV does not account for variability in tissue size and shape. The mean SUV provides insights into the relative intensity of PET signal in each tissue (Fig 14A). This comparison of mean SUV reveals the relative intensity of the signal across tissues, with the prostate having consistently high signal in all animals. In contrast, the total SUV/whole body Total SUV for the different tissues illustrates the percentages of the total whole-body signal in each tissue without consideration of the size of each tissue relative to the other (Fig 14B). All 7 PET parameters for each tissue are shown in the heatmap (Fig 14C), which reveals that different tissues have unique signal characteristics. From this analysis, it is evident that the prostate signal of IN22 and JF82 have the highest mean SUV and standard deviation with relatively a high median SUV being also seen in the prostate of the 3^rd^ animal LP14. For example, the prostate signal of IN22 and JF82 has the highest mean SUV and standard deviation with relatively a high median SUV being also seen in the prostate of the 3^rd^ animal LP14, indicating a more clustered signal compared to the tissues with the highest total SUV.

**Figure 14.**
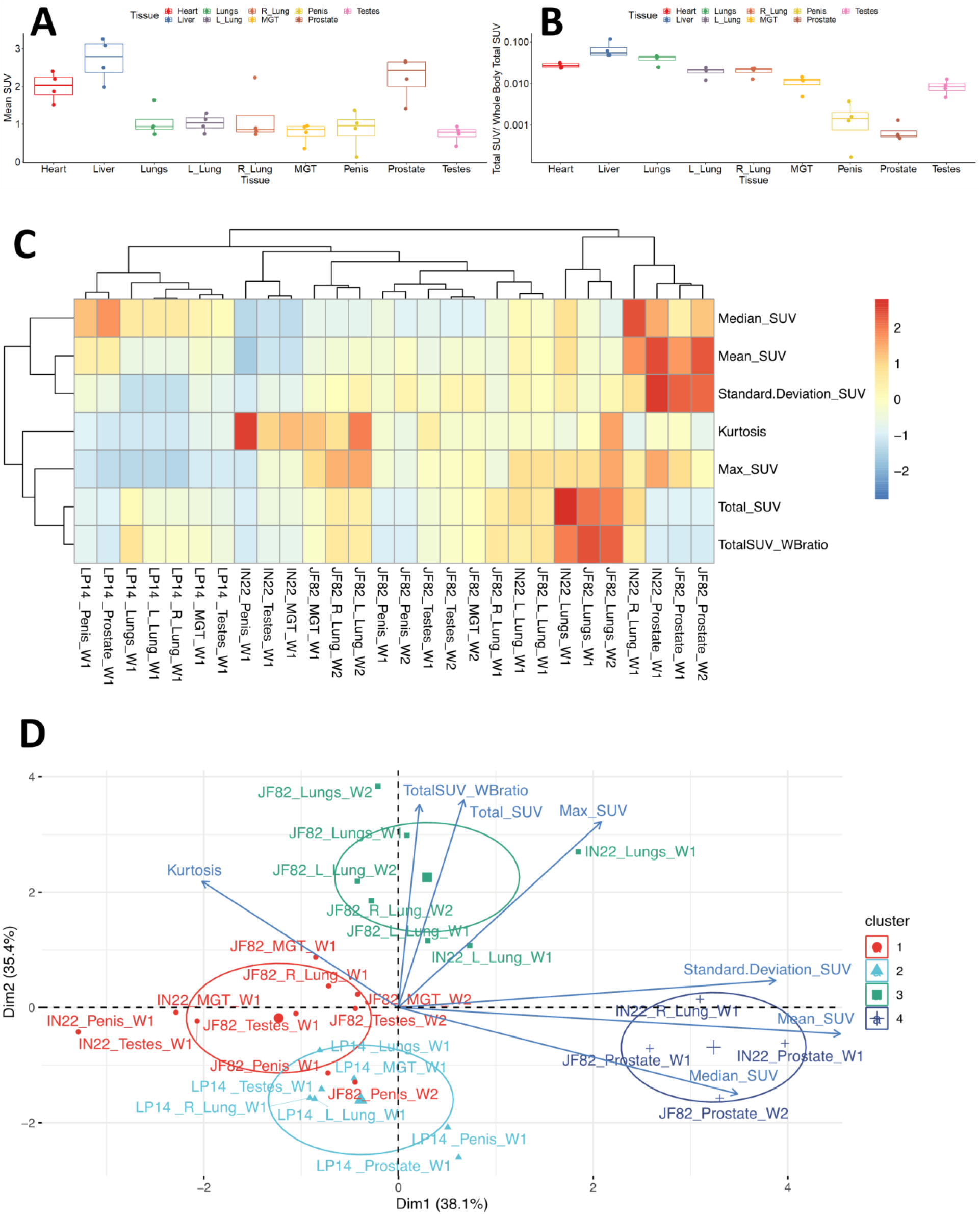
Principle components analysis of SUV measures. (A) Mean SUV values from isolated tissue volumes. (B) The ratio of total SUV for each tissue volume to whole-body total SUV. (C) Heat map showing clustering of tissues and parameters measured from the PET data. (D) Principal components analysis of measures showing hierarchical clustering of tissues and measures.

To facilitate the use of our PET data to gain insights into our data set we utilized Principal Component Analysis (PCA) followed by hierarchical clustering to examine the data from all scans of the 3 animals. To cluster tissues according to their SUV signal characteristics, we applied PCA including all variables obtained from the SUV measurements (i.e., Total SUV, Total SUV Whole Body ratio, Mean SUV, Median SUV, Standard Deviation SUV, Max SUV, and Kurtosis) for each tissue and animal. After this analysis, we observed 4 different clusters that correspond to tissues that display very distinct SUV signals. The clustering captures the prostate signal described in the heat map where they are part of cluster 4 (dark blue group). Of note, the major right lung signal of IN22 also clusters within this group, while the prostate signal of LP14 does not (Fig 14D). The evident PET signal observed in the lungs of the Delta variant animals IN22 and JF82 are part of the cluster 3 (green group) (Fig 14D) revealing their similarity with its small foci of high signal and increased kurtosis value (indicating data heavy tails or more outliers) consistent with the signal variability. The higher kurtosis is also associated with penile and testes signal of all the animals in the red and light blue clusters 1 and 2. This analysis allows us to begin to appreciate the nature of the PET signal associated with the different tissues, different animals, different viruses, and identify outliers.

## Discussion

The goal here was to develop a F(ab’)2 based immunoPET probe to allow the visualization of SARS-CoV-2 spike protein expression in the rhesus macaque infection model. This model is clinically relevant as it was used to develop the highly successful vaccines now available against SARS-CoV-2^31, 32^. The ability to detect anatomical sites of SARS-CoV-2 infection will facilitate our understanding of COVID-19 disease severity and the mechanism of comorbidities in the development of pathology. It could be especially useful in understanding long-COVID and its underlying causes^33, 34^. This pilot study is a key step in the clinical development of this PET probe to visualize SARS-CoV-2 replication and to facilitate treatments for COVID-19. The distribution of the probe is complex, but it can reveal sites of SARS-CoV-2 infection in the rhesus macaque model.

The utility of the probe is best demonstrated by the temporal association of the PET probe with CT detected lung pathology between weeks 1 and 2 in JF82. A PET signal was clearly associated with an opaque region (pneumonia) in the left lung in the week 1 scan, but both the PET signal and CT detected pneumonia are not present in the week 2 scan, validating this technique. The overlays, shown in Fig 9, further illustrate this transient lung pathology detected by comparing the two PET/CT scans separated by one week. Supporting this interpretation of the PET/CT data, the pathology report shows that the left lung of JF82 has type II pneumocyte hyperplasia and inflammatory infiltrate consistent with the normal course of infection observed in the rhesus macaque model. Overall, the lung pathology associated with SARS-CoV-2 in our three animals was consistent with the observation of many infected rhesus macaques where the lung pathology was typically mild and peaked at 1 week after IN/IT challenge. This dynamic was generally observed for lung associated signal in this limited pilot study. There was some consolidation and inflammation observed in animal IN22 and this coincided with an adjacent PET signal in both animals infected with the Delta variant.

Further validation of the PET/CT probe’s ability to identify areas of infection at sites other than the lungs and its utility in understanding COVID-19 pathogenesis was the revelation of a reproducible infection of the MGT. In all three animals, the probe was associated with the prostate, penis, pampiniform plexus, and testicles. We have been able to identify SARS-CoV-2 infected cells in the testicles of all 3 animals. Comparing the longitudinal PET scans of JF82 reveals that while the lung pathology and signal wane between week 1 and 2, the signal of the MGT, and more specifically in the testicles increases in week 2. Consistent with increased testicular infection at week 2 indicated in the PET scan, H&E staining of the JF82 testes revealed a unique pathology consisting of denuded stretches of the twisted and intertwined seminiferous tubules (presenting as a tube cluster). Spermatids were absent in these degenerate regions and the remaining Sertoli cells were undergoing apoptosis, seen through caspase 3 staining (Fig. 10S). Local inflammation, or orchitis, was suggested by immune infiltrates (Fig. 10R). Both the PET signal and pathology is greater in the left testicle of JF82. Staining macrophages and mature spermatids for CD206 and all cells for caspase 3 activation reveals a severe, acute response within short stretches of the seminiferous tubules where the Sertoli cells are undergoing apoptosis due to inflammasome activation (caspase 3 activation), and no spermatids are present consistent with an ongoing acute infection (Fig. 10N-S). We have detected infected cells within the JF82 testes ((Fig. 10T, U) and the relationship between the infected cells and testicular pathology is ongoing. Similar decreases in the cellular content of the seminiferous tubules and sloughing of Sertoli cells and spermatids into the lumen have been reported in multiple studies of autopsy tissues from COVID-19 related fatalities^35–40^.

Our results suggest that SARS-CoV-2 rapidly and efficiently infects multiple tissues of the male genital tract (MGT) early during infection in rhesus macaques. The complex vasculature and known ACE2 expression of the tissues of the MGT make it a potential target of the virus^11, 41–43^. The SARS-CoV-2 infection of the testicles has been reported in mouse and hamster respiratory challenge models^44–46^. Likewise, the testicles are also a target of Ebola and Zika virus during systemic infection^47^. We observed a similar distribution of PET-probe signal in all 3 animals in the week 1 scan with labeling of the prostate, root of the penis, the top the testicles and a second region of labeling at the base of the testicles (Fig. 11E-H). The signal above the testicles localizes to the pampiniform plexus and vasculature of the spermatic cord while the signal at the base of the testes is less clear, but appears to be associated with the cauda epididymis, the highly vascularized tail of the epididymis that serves as the storage site for mature sperm. A further dissection of the testicles before the organ scan should facilitate a detailed localization of the PET signal associated with the MGT in future studies.

Although these studies were done with a rhesus macaque model, it is reasonable to suggest that these observations may also apply to humans infected with SARS-CoV-2 because of several clinical observations relating to male sexual health and fertility. It is highly relevant in this extrapolation to consider that we have identified 4 distinct tissues where SARS-CoV-2 infection could impact male sexual health and fertility: SARS-CoV-2 infection of the prostate, penis, pampiniform plexus, and testicles. The infection of the MGT and associated pathology has been suggested by several publications and clinical studies. The prostate is known to be ACE2 positive.^48^ Interest in SARS-CoV-2 infection of the prostate has focused on two areas. First is the potential impact on treatment of benign prostate hyperplasia^49^ and prostate cancer ^50^ with androgen deprivation therapy on the severity of COVID-19^51^ and secondly, the potential impact of SARS-CoV-2 infection on prostate cancer treatments. It is notable that prostate cancers are known to express high levels of transmembrane serine protease 2, TMPRSS2^52^, which is known to activate the SARS-CoV-2 spike protein to its optimal fusogenic potential^53^. Multiple studies have explored this space and it does not appear that SARS-CoV-2 infection is associated with an increase in prostate cancer^54^. In contrast, another study suggests that infection with SARS-CoV-2 may be associated with an increase in prostate specific antigen (PSA) detection in plasma^55^. Future studies are needed to confirm whether the robust signal of the SARS-CoV-2 PET/CT probe reflects a high-level infection of the human prostate and its subsequent impact on male sexual health and fertility^56, 57^. SARS-CoV-2 infection of the penis is potentially associated with the vasculature of the corpus cavernosum, which expresses high levels of ACE2 in the rhesus macaque and human penile tissue (Fig 7S) ^43, 58^. Because the corpus cavernosum plays a key role in erectile function, the inflammation caused by SARS-CoV-2 infection of the penile vasculature is hypothesized to lead to erectile dysfunction (ED). This has indeed been reported to be linked to COVID-19^43, 59, 60^. In addition, treatments for ED such as Viagra and Cialis are known to affect the renin-angiotensin-aldosterone-system where ACE2 functions as a part of the physiologic regulation of blood flow associated with normal erectile function^61^.

A potential impact of COVID-19 infection on the pampiniform plexus might be suggested by several case reports of COVID-19 associated thrombosis located in the pampiniform plexus^62–65^. Additionally, the signal distribution of the left and right testes is distinct, with the signal of the right testicle being more focused at the top of the testicle while the signal on the left testicle being more distributed in the spermatic cord. This is reminiscent of the condition known as varicocele, which manifests as varicose veins of the scrotum, and is prominent in the left testicle relative to the right testicle^66^. This is due to the left testicle receiving its blood flow from the left renal vein which exposes it to higher blood pressure and slower blood flow^67^. This difference could be insightful if it is confirmed in more animals.

The potential infection of the testicles by SARS-CoV-2 could be highly impactful on male fertility, potentially decreasing sperm count and semen quality^47, 68–70^. It is known that SARS-CoV-2 infection in humans is associated with oligo- and azoospermia and a transient decrease in fertility after infection^36, 38, 66, 68, 71, 72^. One study found that fertility amongst infected men was reduced and returned to baseline 3-6 months after SARS-CoV-2 infection^73^. This decrease in fertility was not seen in infected women or men who received a SARS-CoV-2 vaccination. We find that the pathology associated with the testicles in the week 2 necropsy is extreme, with apparent ablation of sperm production within short regions of the seminiferous tubules and with accompanying immune infiltration consistent with an emerging COVID-19 associated orchitis^64, 74^. Multiple studies have reported a decrease in testosterone after SARS-CoV-2 infection^56, 75, 76^. Leydig cells, which produce testosterone, are known targets of SARS-CoV-2^35, 37, 77^. The decrease in testosterone correlates with disease severity^76, 78^. This susceptibility of the MGT to infection with SARS-CoV-2 may be consideration in the of higher mortality associated with COVID-19 in men compared to women. The direct infection of the testicles observed in the NHP studies presented here are consistent with the MGT pathology reported in men^35–40^. The infection of the MGT may be a common outcome in SARS-CoV-2 infection. We suggest further study in mild, and even in asymptomatic infection, in men and macaques is clearly required. Because of the four distinct mechanisms negatively impacting human male sexual health and fertility, and the observation of all four mechanisms in all animals evaluated to date, we feel compelled to report this information at this early stage of study and evaluation.

Using a novel ^64^Cu-labelled F(ab’)2 probe targeting the SARS-CoV-2 spike protein, we have found that it is possible to identify sites of SARS-CoV-2 infection in the rhesus macaque infection model using a PET/CT scan. The rapid and consistent spread of SARS-CoV-2 infection to the MGT in the first week of infection in the rhesus macaque suggests this could be an important post pandemic health consideration as those infected during the pandemic develop advanced disease and other manifestations of long COVID-19. Currently, there is no other methods that allows for the unbiased discovery of novel anatomical sites of SARS-CoV-2 infection. We believe the immunoPET technique described here will be an important addition to the toolkit for studying and understanding SARS-CoV-2 pathogenesis. The availability of an immunoPET probe to SARS-CoV-2 in the clinical setting has the potential to reveal the underlying role of disseminated viral infection in long COVID and could guide therapeutic interventions to resolve SARS-CoV-2 related sequalae which could be a major health concern for the lifetimes of those infected during the COVID-19 pandemic.

## ACKNOWLEDGEMENTS

This work was supported by a NOSI supplement (R37AI094595-10S1 PI: Hope) to parent grant R37AI094595 (PI Hope) and by NIH grants R21AI163912 (PI Hultquist) P51OD01110459, and U19AI135964 (E.A.O.). We would like to thank Nicholas Maness for supplying the delta variant virus and Maury Duplantis, Brooke F. Gasperge, Kasi E. Russell-Lodrigue, and the Veterinary team at TNPRC. Thanks to Rich D’Aquila for critical reading and editing of the manuscript.

## Material and Methods

### Preparation of F(ab)2

CR3022-IgG1 was acquired from Absolute Antibody (#Ab01680-10.0, Absolute Antibody). The IgG1 was digested into F(ab)2 as previously described. Briefly, a Genovis FragIT antibody digestion kit (#A2-FR2-100, Genovis) was used according to manufacturer’s protocol. CR3022 was digested by adding to the FragIT column and rocking for 1 hr. at room temperature. The column was then centrifuged to elute fragments. Fc fragments were removed after incubation with the CaptureSelect Fc Column. F(ab’)2 fragments were eluted from this column and size was confirmed using SDS-PAGE.

### DOTA Labeling of Antibody

CR3022-F(ab)2 was labeled with the chelator dodecane tetra-acetic acid (DOTA) for the attachment of ^64^Cu. Chelex 100 chelating resin (#142-1253, BioRad) was used to prepare two buffers: 0.1M sodium phosphate (pH 7.3) and 0.1M ammonium acetate (pH 5.5). Five grams of Chelex was added to 100ml of each buffer and stirred at room temperature for 1 hour. Buffers were then filter sterilized using 0.22 µM filters. CR3022-F(ab’)2 was buffer exchanged into the prepared 0.1 M sodium phosphate using Zeba columns (ThermoFisher, USA). DOTA-NHS-ester (#B-280, Macrocyclics) was dissolved in dimethyl sulfoxide (DMSO) at a concentration of 10mM. A 1:10 dilution of 1M sodium bicarbonate was made into a tube containing F(ab’)2 in sodium phosphate and 10mM DOTA was added at a molar ratio of 5:1. The tube was mixed and left to rock at 37C for 1.5 hours. The labeled F(ab’)2 was then buffer exchanged into fresh 0.1 M sodium phosphate using a Zeba column.

### ^64^Cu labeling

Before labeling DOTA conjugated F(ab’)2 was buffer exchanged into freshly prepared 0.1M ammonium acetate (pH 5.5) prepared with Chelex 100 using 10kDa cutoff Amicon centrifugal microfilters (Cork, IRL). Cu^64^ chloride was obtained from Washington University, St. Louis MO and shipped overnight to TNPRC. All work with Cu^64^ was performed inside a lead castle. The radioactivity of the Cu^64^ was measured using an Atom Force 500 dose calibrator and recorded. Next the Cu^64^ was added to a tube containing the DOTA labeled F(ab’)2 in 0.1 M ammonium acetate and incubated at 37°C for 1 hour while rotating. The labeled F(ab’)2 probe was then separated from unlabeled Cu^64^ and buffer exchanged into PBS via sequential centrifugation steps using 10kDa cutoff Amicon centrifugal microfilters to a final volume of ∼20µl. The conjugated probe was then diluted to the final volume in PBS in sterile glass vials and drawn into single dose syringes. Each dose typically labelled in the 1-3 mCi per dose.

### Animals and virus

The virus used for experimental infection of LP14 was SARS-CoV-2; 2019-nCoV/USA-WA1/2020 (https://www.ncbi.nlm.nih.gov/nuccore; accession number MN985325.1) obtained from BEI resources (BEI #NR-52281). Virus stock was prepared in Vero E6 cells and sequence confirmed by deep sequencing. Plaque assays were performed in Vero E6 cells. Vero E6 cells were acquired from ATCC (Manassas, VA). The virus used for experimental infection of IN22 and JF82 was SARS-CoV-2; hCoV-19/USA/MD-HP05647/2021 (Lineage B.1.617.2; Delta variant) obtained from BEI resources (BEI #NR-55672). Delta variant stock was expanded using Calu-3 cells and prepared as above using Vero E6 cells.

Three male rhesus macaques (Macaca mulatta) were used in this study. All animals were housed at the Tulane National Primate Research Center (TNPRC, Covington, LA) which is accredited by the Association for the Assessment and Accreditation of Laboratory Animal Care. All procedures were reviewed and approved by the Tulane University Institutional Animal Use Committee under protocol number P0452.One animal was inoculated with 1.1×10^6^ tissue culture infectious dose (TCID_50_) of the WA1 strain of SARS-CoV-2 via concomitant intratracheal/intranasal instillation (1mL intratracheal, 500µL per each nare). This animal was necropsied 8 days post-infection. The remaining two animals were inoculated with 1.12×10^6^ TCID_50_ of the delta variant of SARS-CoV-2 via the same route. One animal was necropsied at day 7 and one at day 14 post-infection. All animals were intravenously infused with 0.5 mg/kg of the radiolabeled F(ab’)2 probe and JF82 was infused on two separate occasions. Macaque LP14 received 4 mCi, JF82 and IN22 received 1.3 and 1,4 mCi respectively on Dec 9, and JF82 received an additional dose of 0.7 mCi on Dec 15. LP14 had nasal and salivary swabs taken at day 7. IN22 had swabs taken at days 3, 5, and 7 post-infection, while JF82 had swabs taken at days 3, 5, 7, and 14.

### PET/CT Imaging and Necropsy

PET/CT guided necropsies were performed in three sequential phases each separated by a PET/CT followed by a period of analysis and sampling (1. whole-body, 2. organ, and 3. tissue). First, whole-body scans were acquired prior to sending the animal to necropsy. Following euthanasia, all major organ systems were removed (pluck, gastrointestinal tract, liver, spleen, kidneys, urinary bladder, testicles, penis, prostate, seminal vesicles, nasal turbinate, lymph nodes, carotid artery, cervical spinal cord, and brain) placed in a clear, plastic, sealable container and sent back to PET/CT for an “organ scan”. After the organ scans were reconstructed, “hot” regions of each major organ (as seen on the organ scan) were sampled and placed in cryomolds. The final “tissue” scan was acquired by placing the cryomolds in a clear, plastic, sealable container and scanning them with the PET/CT. Following acquisition of the tissue scan, cryomolds were filled with OCT and frozen on dry ice. All remaining tissue (not placed in cryomolds) was placed in zinc-formalin fixative. All samples were stored for 5 days before being removed from containment.

### PET/CT Image Analysis

Acquired PET/CT whole-body images were analyzed using the MIM Software (MIM Software Inc., Cleveland, OH). The PET and CT scans were reconstructed using the software and PETCT fusions were created to analyze regions of interest through axial, sagittal, and coronal views. The PET scans were presented in calculated Standardized Uptake Values, and all images were set to the same scale (0-20 SUVbw). The PET scale was selected based on the overall signal intensity of the PET scans, and the CT scale for optimal visibility of the tissues. Regions of interest (ROI) were isolated using a combination of the Region Grow function and manual contouring on a representative scan, then these regions were copied on subsequent scans of the same animal using a specialized developed workflow. This workflow allows the software to use the CT scans to map the selected ROI and locate that exact volume in subsequent scans. Manual adjustments were then used to counter any changes in the animals’ orientation between scans. The areas within these regions were then extracted from the full scans, and the anatomical regions were analyzed in both 2D cross-sections and 3D projections. 3D views are maximum intensity projections of isolated ROI in the PET scans.

To cluster tissues according to their SUV signal characteristics, we used Principal Component Analysis (PCA) where we included all variables obtained from the SUV measurements (i.e., Total SUV, Total SUV Whole Body ratio, Mean SUV, Median SUV, Standard Deviation SUV, Max SUV, and Kurtosis) for each tissue and animal. The clustering of each tissue was subsequently obtained by agglomerative hierarchical clustering on the PCA results. After building an initial hierarchical tree, the sum of within-cluster inertia was calculated for each partition. The selected partition was the one with the higher relative loss of inertia. Both PCA and agglomerative hierarchical clustering were performed using FactoMineR package and Factoextra was used for visualization of the clustering results.

### Isolation and Quantification of viral RNA

Viral load was quantified from pharyngeal and nasal swabs taken from infected animals at TNPRC as previously described (1) Swab and bronchial brush samples were collected in 200 μL of DNA/RNA Shield 1× (catalog number R1200; Zymo Research, Irvine, CA) and extracted for viral RNA using the Quick-RNA viral kit (catalog number R1034/5; Zymo Research). The Viral RNA Buffer was dispensed directly to the swab in the DNA/RNA Shield. A modification to the manufacturers’ protocol was made to insert the swab directly into the spin column to centrifugate, allowing all the solution to cross the spin column membrane. The viral RNA was then eluted (45 μL) from which 5 μL was added in a 0.1-mL fast 96-well optical microtiter plate format (catalog number 4346906; Thermo Fisher Scientific, Waltham, MA) for a 20-μL real-time quantitative RT-PCR (RT-qPCR) reaction. The RT-qPCR reaction used TaqPath 1-Step Multiplex Master Mix (catalog number A28527; Thermo Fisher Scientific) along with the 2019-nCoV RUO Kit (catalog number 10006713; IDTDNA, Coralville, IA), a premix of forward and reverse primers and a FAM-labeled probe targeting the N1 amplicon of the N gene of SARS2-nCoV19 (https://www.ncbi.nlm.nih.gov/nuccore; accession number MN908947). The reaction master mix was added using an X-stream repeating pipette (Eppendorf, Hauppauge, NY) to the microtiter plates, which were covered with optical film (catalog number 4311971; Thermo Fisher Scientific), vortexed, and pulse centrifuged. The RT-qPCR reaction was subjected to RT-qPCR at a program of uracil-DNA glycosylase incubation at 25°C for 2 minutes, room temperature incubation at 50°C for 15 minutes, and an enzyme activation at 95°C for 2 minutes followed by 40 cycles of a denaturing step at 95°C for 3 seconds and annealing at 60°C for 30 seconds. Fluorescence signals were detected with a QuantStudio 6 Sequence Detector (Applied Biosystems, Foster City, CA). Data were captured and analyzed with Sequence Detector Software version 1.3 (Applied Biosystems). Viral copy numbers were calculated by plotting Cq values obtained from unknown (i.e., test) samples against a standard curve that represented known viral copy numbers. The limit of detection of the viral RNA assay was 10 copies per reaction volume. A 2019-nCoV–positive control (catalog number 10006625; IDTDNA) was analyzed in parallel with every set of test samples to verify that the RT-qPCR master mix and reagents were prepared correctly to produce amplification of the target nucleic acid. A non-template control was included in the qPCR to ensure that there was no cross-contamination between reactions.

OCT embedded tissue blocks were sectioned between 10-15 µM and 2-4 sections were placed into RNAse free tubes. RNA extraction was carried out using a RNeasy Plus Mini Kit (#74124, Qiagen), according to manufacturer’s protocol. Briefly, 350ul of lysis buffer with dithiothreitol (DTT) was added to the tubes containing sections. The tubes were vortexed briefly then frozen at −20C. Once thawed the samples were again vigorously vortexed then centrifuged for 3 minutes. Supernatant was removed and applied to the gDNA Eliminator spin column. The flow-through was mixed with 350ul of 70% ethanol and added to a RNeasy spin column. The spin column was washed three times with buffers RW1 and RPE. RNA was then eluted using 30-50ul of RNAse free water. All steps prior to addition of lysis buffer were carried out in a BSL3 facility.

### Tissue Processing and H&E Staining

Tissue samples were collected in Zinc formalin (Anatech) and fixed for a minimum of 72 hours before being washed and dehydrated using a Thermo Excelsior AS processor. Upon removal from the processor, tissues were transferred to a Thermo Shandon Histocentre 3 embedding station where they were submersed in warm paraffin and allowed to cool into blocks. From these blocks, 5um sections were cut and mounted on charged glass slides, baked overnight at 60°C, and passed through Xylene, graded ethanol, and double distilled water to remove paraffin and rehydrate tissue sections. A Leica Autostainer XL was used to complete the deparaffinization, rehydration and routine hematoxylin and eosin stain. Slides were digitally imaged with a NanoZoomer S360 (Hamamatsu) and subsequently examined by a board-certified, veterinary pathologist using HALO software (Indica Labs).

### Fluorescent Immunohistochemistry of FFPE tissues

5um sections of Formalin-fixed, paraffin-embedded (FFPE) tissues were mounted on charged glass slides, baked for two hours at 60°C, and passed through Xylene, graded ethanol, and double distilled water to remove paraffin and rehydrate tissue sections. A microwave was used for heat induced epitope retrieval. Slides were heated in a high pH solution (Vector Labs H-3301), rinsed in hot water, and transferred to a heated low pH solution (Vector Labs H-3300) where they were allowed to cool to room temperature. Sections were washed in a solution of phosphate-buffered saline and fish gelatin (PBS-FSG) and transferred to a humidified chamber, for staining at room temperature. Lung sections were blocked with 10% normal goat serum (NGS) for 40 minutes, followed by a 60-minute incubation with the anti-SARS primary antibody diluted in NGS. Slides were washed and transferred back to the humidified chamber for a 40-minute incubation with the secondary antibody, also diluted in NGS. Sequential staining of FFPE testes, for CD206 and Caspase 3, was done as described above with 1% normal donkey serum (NDS) being used in place of NGS for blocking and antibody dilutions. DAPI was used to label the nuclei of each section. Slides were mounted using a homemade anti-quenching mounting media containing Mowiol (Calbiochem #475904) and DABCO (Sigma #D2522) and imaged at 20X with a Zeiss Axio Slide Scanner.

| PRIMARY ANTIBODY | SPECIES | COMPANY | CATALOG | DILUTION | SECONDARY ANTIBODY |
| --- | --- | --- | --- | --- | --- |
| SARS | Guinea Pig | BEI | NR-10361 | 1:1000 | Goat anti-guinea pig Alexa Fluor488 |
| CD206 | Goat | R&D Systems | AF2532 | 1:50 | Donkey anti-goat Alexa Fluor488 |
| Caspase 3 | Rabbit | Abcam | Ab13847 | 1:200 | Donkey anti-rabbit Alexa Fluor555 |
| DAPI |  | Invitrogen | D1306 | 1:20,000 |  |

### Fluorescence Microscopy of OCT embedded tissues

OCT embedded tissue were cryosectioned in a BSL3 facility between 10 and 15 µM. One or two sections of each tissue were placed on glass microscope slides. Tissues were placed into an airtight container containing 4% PFA in PIPES buffer. The container was sealed and thoroughly decontaminated before being removed from the BSL3 facility. The remainder of the staining took place outside of the BSL3. Tissue sections were treated with L-lysine (0.1%, SigmaAldrich) to reduce background and non-specific interactions before being blocked using 3% BSA (Invitrogen, ThermoFisher) for 30 minutes at room temperature. Staining with primary antibodies was carried out at 37°C and secondary antibodies at room temperature. each for 1hr. Table of primary and secondary antibodies used is below. All slides were stained with Hoechst (1:25,000, ThermoFisher, USA) for 15 minutes and washed with PBS between all steps. Dako fluorescent mounting medium (#S302380-2, Agilent, USA) was used to mount cover slips which were sealed with nail polish. A DeltaVision Ultra inverted microscope (Cytivia, USA) was used to obtain images using the 10x, 20x, 60x, and 100x lenses. Images were deconvolved, stitched, and projected using the softWoRx software (Applied Precision, USA).

| Clone | Antibody | Target | Company | Catalog# | Dilution |
| --- | --- | --- | --- | --- | --- |
|  | Rb pAb | Spike | Abcam | Ab272504 | 1:200 |
| 3A2 | Ms mAb | Spike | Abcam | Ab272420 | 1:200 |
| 6H3 | Ms mAb | NC | GeneTex | GTX632269 | 1:200 |
| 5A10 | Ms mAb | NSP8 | GeneTex | GTX632696 | 1:200 |
|  | Rb pAb | NC | Novus Biologicals | NB100-56576 | 1:200 |
|  | Rb pAb | NC | GeneTex | GTX135357 | 1:200 |
| rJ2 | Ms mAb j2 | Anti-double stranded RNA | Millipore Sigma | MABE1134 | 1:1000 |

## REFERENCES

1. Cheung CCL, Goh D, Lim X, Tien TZ, Lim JCT, Lee JN, Tan B, Tay ZEA, Wan WY, Chen EX, Nerurkar SN, Loong S, Cheow PC, Chan CY, Koh YX, Tan TT, Kalimuddin S, Tai WMD, Ng JL, Low JG-H, Yeong J, Lim KH. Residual SARS-CoV-2 viral antigens detected in GI and hepatic tissues from five recovered patients with COVID-19. Gut. 2021. doi: 10.1136/GUTJNL-2021-324280.

2. Kariyawasam JC, Jayarajah U, Riza R, Abeysuriya V, Seneviratne SL. Gastrointestinal manifestations in COVID-19. Trans R Soc Trop Med Hyg. 2021;115(12):1362–88. doi: 10.1093/trstmh/trab042. PubMed PMID: 33728439; PMCID: PMC7989191.

3. Keshavarz P, Rafiee F, Kavandi H, Goudarzi S, Heidari F, Gholamrezanezhad A. Ischemic gastrointestinal complications of COVID-19: a systematic review on imaging presentation. Clin Imaging. 2021;73:86–95. Epub 20201208. doi: 10.1016/j.clinimag.2020.11.054. PubMed PMID: 33341452; PMCID: PMC7837247.

4. Javadekar NS. Gut in COVID 19-is it worth noticing. Ind Psychiatry J. 2021;30(Suppl 1):S267–S9. Epub 20211022. doi: 10.4103/0972-6748.328826. PubMed PMID: 34908706; PMCID: PMC8611584.

5. Paterson BJ, Durrheim DN. Wastewater surveillance: an effective and adaptable surveillance tool in settings with a low prevalence of COVID-19. Lancet Planet Health. 2022;6(2):e87–e8. doi: 10.1016/S2542-5196(22)00009-2. PubMed PMID: 35150632; PMCID: PMC8828367.

6. Gregory DA, Wieberg CG, Wenzel J, Lin CH, Johnson MC. Monitoring SARS-CoV-2 Populations in Wastewater by Amplicon Sequencing and Using the Novel Program SAM Refiner. Viruses. 2021;13(8). Epub 20210819. doi: 10.3390/v13081647. PubMed PMID: 34452511; PMCID: PMC8402658.

7. Nakamura Y, Katano H, Nakajima N, Sato Y, Suzuki T, Sekizuka T, Kuroda M, Izutani Y, Morimoto S, Maruyama J, Koie M, Kitamura T, Ishikura H. SARS-CoV-2 is localized in cardiomyocytes: a postmortem biopsy case. Int J Infect Dis. 2021;111:43–6. Epub 20210809. doi: 10.1016/j.ijid.2021.08.015. PubMed PMID: 34384897; PMCID: PMC8351278.

8. Lei J, Liu Y, Xie T, Yao G, Wang G, Diao B, Song J. Evidence for residual SARS-CoV-2 in glioblastoma tissue of a convalescent patient. Neuroreport. 2021;32(9):771–5. doi: 10.1097/WNR.0000000000001654. PubMed PMID: 33994523.

9. Song E, Zhang C, Israelow B, Lu-Culligan A, Prado AV, Skriabine S, Lu P, Weizman OE, Liu F, Dai Y, Szigeti-Buck K, Yasumoto Y, Wang G, Castaldi C, Heltke J, Ng E, Wheeler J, Alfajaro MM, Levavasseur E, Fontes B, Ravindra NG, Van Dijk D, Mane S, Gunel M, Ring A, Kazmi SAJ, Zhang K, Wilen CB, Horvath TL, Plu I, Haik S, Thomas JL, Louvi A, Farhadian SF, Huttner A, Seilhean D, Renier N, Bilguvar K, Iwasaki A. Neuroinvasion of SARS-CoV-2 in human and mouse brain. J Exp Med. 2021;218(3). doi: 10.1084/jem.20202135. PubMed PMID: 33433624; PMCID: PMC7808299.

10. Hanley B, Naresh KN, Roufosse C, Nicholson AG, Weir J, Cooke GS, Thursz M, Manousou P, Corbett R, Goldin R, Al-Sarraj S, Abdolrasouli A, Swann OC, Baillon L, Penn R, Barclay WS, Viola P, Osborn M. Histopathological findings and viral tropism in UK patients with severe fatal COVID-19: a post-mortem study. Lancet Microbe. 2020;1(6):e245–e53. Epub 20200820. doi: 10.1016/S2666-5247(20)30115-4. PubMed PMID: 32844161; PMCID: PMC7440861.

11. Edenfield RC, Easley CAt. Implications of testicular ACE2 and the renin-angiotensin system for SARS-CoV-2 on testis function. Nat Rev Urol. 2022;19(2):116–27. Epub 20211126. doi: 10.1038/s41585-021-00542-5. PubMed PMID: 34837081; PMCID: PMC8622117.

12. Collins AB, Zhao L, Zhu Z, Givens NT, Bai Q, Wakefield MR, Fang Y. Impact of COVID-19 on Male Fertility. Urology. 2022. doi: 10.1016/J.UROLOGY.2021.12.025.

13. Parker AM, Brigham E, Connolly B, McPeake J, Agranovich AV, Kenes MT, Casey K, Reynolds C, Schmidt KFR, Kim SY, Kaplin A, Sevin CM, Brodsky MB, Turnbull AE. Addressing the post-acute sequelae of SARS-CoV-2 infection: a multidisciplinary model of care. Lancet Respir Med. 2021;9(11):1328–41. Epub 20211019. doi: 10.1016/S2213-2600(21)00385-4. PubMed PMID: 34678213; PMCID: PMC8525917.

14. Zhao JM, Zhou GD, Sun YL, Wang SS, Yang JF, Meng EH, Pan D, Li WS, Zhou XS, Wang YD, Lu JY, Li N, Wang DW, Zhou BC, Zhang TH. [Clinical pathology and pathogenesis of severe acute respiratory syndrome]. Zhonghua Shi Yan He Lin Chuang Bing Du Xue Za Zhi. 2003;17(3):217–21. PubMed PMID: 15340561.

15. Xu J, Qi L, Chi X, Yang J, Wei X, Gong E, Peh S, Gu J. Orchitis: a complication of severe acute respiratory syndrome (SARS). Biol Reprod. 2006;74(2):410–6. Epub 20051019. doi: 10.1095/biolreprod.105.044776. PubMed PMID: 16237152; PMCID: PMC7109827.

16. Beddingfield BJ, Maness NJ, Fears AC, Rappaport J, Aye PP, Russell-Lodrigue K, Doyle-Meyers LA, Blair RV, Carias AM, Madden PJ, Redondo RL, Gao H, Montefiori D, Hope TJ, Roy CJ. Effective prophylaxis of COVID-19 in rhesus macaques using a combination of two parentally-administered SARS-CoV-2 neutralizing antibodies. bioRxiv. 2021:2021.05.26.445878-2021.05.26. doi: 10.1101/2021.05.26.445878.

17. Blair RV, Vaccari M, Doyle-Meyers LA, Roy CJ, Russell-Lodrigue K, Fahlberg M, Monjure CJ, Beddingfield B, Plante KS, Plante JA, Weaver SC, Qin X, Midkiff CC, Lehmicke G, Golden N, Threeton B, Penney T, Allers C, Barnes MB, Pattison M, Datta PK, Maness NJ, Birnbaum A, Fischer T, Bohm RP, Rappaport J. Acute Respiratory Distress in Aged, SARS-CoV-2–Infected African Green Monkeys but Not Rhesus Macaques. The American Journal of Pathology. 2021;191(2):274–82. doi: 10.1016/J.AJPATH.2020.10.016.

18. Chandrashekar A, Liu J, Martino AJ, McMahan K, Mercad NB, Peter L, Tostanosk LH, Yu J, Maliga Z, Nekorchuk M, Busman-Sahay K, Terry M, Wriji LM, Ducat S, Martine DR, Atyeo C, Fischinger S, Burk JS, Slei MD, Pessaint L, Van Ry A, Greenhouse J, Taylor T, Blade K, Cook A, Finneyfrock B, Brown R, Teow E, Velasco J, Zahn R, Wegmann F, Abbink P, Bondzi EA, Dagotto G, Gebr MS, He X, Jacob-Dolan C, Kordana N, Li Z, Lifto MA, Mahrokhia SH, Maxfiel LF, Nityanandam R, Nkolol JP, Schmid AG, Mille AD, Bari RS, Alter G, Sorge PK, Este JD, Andersen H, Lewi MG, Barou DH. SARS-CoV-2 infection protects against rechallenge in rhesus macaques. Science. 2020;369(6505):812–7. doi: 10.1126/SCIENCE.ABC4776.

19. Fahlberg MD, Blair RV, Doyle-Meyers LA, Midkiff CC, Zenere G, Russell-Lodrigue KE, Monjure CJ, Haupt EH, Penney TP, Lehmicke G, Threeton BM, Golden N, Datta PK, Roy CJ, Bohm RP, Maness NJ, Fischer T, Rappaport J, Vaccari M. Cellular events of acute, resolving or progressive COVID-19 in SARS-CoV-2 infected non-human primates. Nat Commun. 2020;11(1):6078. Epub 20201127. doi: 10.1038/s41467-020-19967-4. PubMed PMID: 33247138; PMCID: PMC7695721.

20. Blair RV, Vaccari M, Doyle-Meyers LA, Roy CJ, Russell-Lodrigue K, Fahlberg M, Monjure CJ, Beddingfield B, Plante KS, Plante JA, Weaver SC, Qin X, Midkiff CC, Lehmicke G, Golden N, Threeton B, Penney T, Allers C, Barnes MB, Pattison M, Datta PK, Maness NJ, Birnbaum A, Fischer T, Bohm RP, Rappaport J. Acute Respiratory Distress in Aged, SARS-CoV-2-Infected African Green Monkeys but Not Rhesus Macaques. Am J Pathol. 2021;191(2):274–82. Epub 20201107. doi: 10.1016/j.ajpath.2020.10.016. PubMed PMID: 33171111; PMCID: PMC7648506.

21. Santangelo PJ, Rogers KA, Zurla C, Blanchard EL, Gumber S, Strait K, Connor-Stroud F, Schuster DM, Amancha PK, Hong JJ, Byrareddy SN, Hoxie JA, Vidakovic B, Ansari AA, Hunter E, Villinger F. Whole-body immunoPET reveals active SIV dynamics in viremic and antiretroviral therapy-treated macaques. Nat Methods. 2015;12(5):427–32. Epub 2015/03/10. doi: 10.1038/nmeth.3320. PubMed PMID: 25751144; PMCID: PMC4425449.

22. Taylor RA, McRaven MD, Carias AM, Anderson MR, Matias E, Arainga M, Allen EJ, Rogers KA, Gupta S, Kulkarni V, Lakhashe S, Lorenzo-Redondo R, Thomas Y, Strickland A, Villinger FJ, Ruprecht RM, Hope TJ. Localization of infection in neonatal rhesus macaques after oral viral challenge. PLoS Pathog. 2021;17(11):e1009855. Epub 20211118. doi: 10.1371/journal.ppat.1009855. PubMed PMID: 34793582; PMCID: PMC8639050.

23. Taylor RA, Xiao S, Carias AM, McRaven MD, Thakkar DN, Arainga M, Allen EJ, Rogers KA, Kumarapperuma SC, Gong S, Fought AJ, Anderson MR, Thomas Y, Schneider JR, Goins B, Fox P, Villinger FJ, Ruprecht RM, Hope TJ. PET/CT targeted tissue sampling reveals virus specific dIgA can alter the distribution and localization of HIV after rectal exposure. PLoS Pathog. 2021;17(6):e1009632. Epub 20210601. doi: 10.1371/journal.ppat.1009632. PubMed PMID: 34061907; PMCID: PMC8195437.

24. Madden PJ, Arif MS, Becker ME, McRaven MD, Carias AM, Lorenzo-Redondo R, Xiao S, Midkiff CC, Blair RV, Potter EL, Martin-Sancho L, Dodson A, Martinelli E, Todd JM, Villinger FJ, Chanda SK, Aye PP, Roy CJ, Roederer M, Lewis MG, Veazey RS, Hope TJ. Development of an In Vivo Probe to Track SARS-CoV-2 Infection in Rhesus Macaques. Front Immunol. 2021;12:810047. Epub 20211224. doi: 10.3389/fimmu.2021.810047. PubMed PMID: 35003140; PMCID: PMC8739270.

25. Fahlberg MD, Blair RV, Doyle-Meyers LA, Midkiff CC, Zenere G, Russell-Lodrigue KE, Monjure CJ, Haupt EH, Penney TP, Lehmicke G, Threeton BM, Golden N, Datta PK, Roy CJ, Bohm RP, Maness NJ, Fischer T, Rappaport J, Vaccari M. Cellular events of acute, resolving or progressive COVID-19 in SARS-CoV-2 infected non-human primates. Nature Communications 2020 11:1. 2020;11(1):1-14. doi: 10.1038/s41467-020-19967-4.

26. Gao G, Hu X, Zhou Y, Rao J, Zhang X, Peng Y, Zhao J, Yao Y, Liu K, Liang M, Liu H, Deng F, Xia H, Shan C, Yuan Z. Infection and pathogenesis of the Delta variant of SARS-CoV-2 in Rhesus macaque. Virologica Sinica. 2022. doi: 10.1016/j.virs.2022.02.001.

27. Nicolas N, Michel V, Bhushan S, Wahle E, Hayward S, Ludlow H, de Kretser DM, Loveland KL, Schuppe HC, Meinhardt A, Hedger MP, Fijak M. Testicular activin and follistatin levels are elevated during the course of experimental autoimmune epididymo-orchitis in mice. Sci Rep. 2017;7:42391. Epub 20170213. doi: 10.1038/srep42391. PubMed PMID: 28205525; PMCID: PMC5304336.

28. Cardona-Maya W, Velilla PA, Montoya CJ, Cadavid A, Rugeles MT. In vitro human immunodeficiency virus and sperm cell interaction mediated by the mannose receptor. J Reprod Immunol. 2011;92(1-2):1–7. Epub 20111019. doi: 10.1016/j.jri.2011.09.002. PubMed PMID: 22015004.

29. Cardona-Maya W, Lopez-Herrera A, Velilla-Hernandez P, Rugeles MT, Cadavid AP. The role of mannose receptor on HIV-1 entry into human spermatozoa. Am J Reprod Immunol. 2006;55(4):241–5. doi: 10.1111/j.1600-0897.2005.00340.x. PubMed PMID: 16533334.

30. Benoff S, Hurley I, Cooper GW, Mandel FS, Hershlag A, Scholl GM, Rosenfeld DL. Fertilization potential in vitro is correlated with head-specific mannose-ligand receptor expression, acrosome status and membrane cholesterol content. Hum Reprod. 1993;8(12):2155–66. doi: 10.1093/oxfordjournals.humrep.a137997. PubMed PMID: 8150918.

31. Corbett KS, Flynn B, Foulds KE, Francica JR, Boyoglu-Barnum S, Werner AP, Flach B, O’Connell S, Bock KW, Minai M, Nagata BM, Andersen H, Martinez DR, Noe AT, Douek N, Donaldson MM, Nji NN, Alvarado GS, Edwards DK, Flebbe DR, Lamb E, Doria-Rose NA, Lin BC, Louder MK, O’Dell S, Schmidt SD, Phung E, Chang LA, Yap C, Todd J-PM, Pessaint L, Ry AV, Browne S, Greenhouse J, Putman-Taylor T, Strasbaugh A, Campbell T-A, Cook A, Dodson A, Steingrebe K, Shi W, Zhang Y, Abiona OM, Wang L, Pegu A, Yang ES, Leung K, Zhou T, Teng IT, Widge A, Gordon I, Novik L, Gillespie RA, Loomis RJ, Moliva JI, Stewart-Jones G, Himansu S, Kong W-P, Nason MC, Morabito KM, Ruckwardt TJ, Ledgerwood JE, Gaudinski MR, Kwong PD, Mascola JR, Carfi A, Lewis MG, Baric RS, McDermott A, Moore IN, Sullivan NJ, Roederer M, Seder RA, Graham BS. Evaluation of the mRNA-1273 Vaccine against SARS-CoV-2 in Nonhuman Primates. https://doiorg/101056/NEJMoa2024671. 2020;383(16):1544–55. doi: 10.1056/NEJMOA2024671.

32. Lindsay KE, Bhosle SM, Zurla C, Beyersdorf J, Rogers KA, Vanover D, Xiao P, Araínga M, Shirreff LM, Pitard B, Baumhof P, Villinger F, Santangelo PJ. Visualization of early events in mRNA vaccine delivery in non-human primates via PET–CT and near-infrared imaging. Nature Biomedical Engineering 2019 3:5. 2019;3(5):371–80. doi: 10.1038/s41551-019-0378-3.

33. Soriano JB, Murthy S, Marshall JC, Relan P, Diaz JV, Condition WHOCCDWGoP-C-. A clinical case definition of post-COVID-19 condition by a Delphi consensus. Lancet Infect Dis. 2021. Epub 20211221. doi: 10.1016/S1473-3099(21)00703-9. PubMed PMID: 34951953; PMCID: PMC8691845.

34. Ward H, Flower B, Garcia PJ, Ong SWX, Altmann DM, Delaney B, Smith N, Elliott P, Cooke G. Global surveillance, research, and collaboration needed to improve understanding and management of long COVID. Lancet. 2021;398(10316):2057–9. Epub 20211110. doi: 10.1016/S0140-6736(21)02444-2. PubMed PMID: 34774190; PMCID: PMC8580495.

35. Duarte-Neto AN, Teixeira TA, Caldini EG, Kanamura CT, Gomes-Gouvea MS, Dos Santos ABG, Monteiro RAA, Pinho JRR, Mauad T, da Silva LFF, Saldiva PHN, Dolhnikoff M, Leite KRM, Hallak J. Testicular pathology in fatal COVID-19: A descriptive autopsy study. Andrology. 2022;10(1):13–23. Epub 20210716. doi: 10.1111/andr.13073. PubMed PMID: 34196475; PMCID: PMC8444746.

36. Moghimi N, Eslami Farsani B, Ghadipasha M, Mahmoudiasl GR, Piryaei A, Aliaghaei A, Abdi S, Abbaszadeh HA, Abdollahifar MA, Forozesh M. COVID-19 disrupts spermatogenesis through the oxidative stress pathway following induction of apoptosis. Apoptosis. 2021;26(7-8):415–30. Epub 20210602. doi: 10.1007/s10495-021-01680-2. PubMed PMID: 34076792; PMCID: PMC8170653.

37. Yang M, Chen S, Huang B, Zhong JM, Su H, Chen YJ, Cao Q, Ma L, He J, Li XF, Li X, Zhou JJ, Fan J, Luo DJ, Chang XN, Arkun K, Zhou M, Nie X. Pathological Findings in the Testes of COVID-19 Patients: Clinical Implications. Eur Urol Focus. 2020;6(5):1124–9. Epub 20200531. doi: 10.1016/j.euf.2020.05.009. PubMed PMID: 32563676; PMCID: PMC7261470.

38. Peirouvi T, Aliaghaei A, Eslami Farsani B, Ziaeipour S, Ebrahimi V, Forozesh M, Ghadipasha M, Mahmoudiasl GR, Aryan A, Moghimi N, Abdi S, Raoofi A, Kargar Godaneh M, Abdollahifar MA. COVID-19 disrupts the blood-testis barrier through the induction of inflammatory cytokines and disruption of junctional proteins. Inflamm Res. 2021;70(10-12):1165–75. Epub 20210826. doi: 10.1007/s00011-021-01497-4. PubMed PMID: 34436630; PMCID: PMC8387554.

39. Achua JK, Chu KY, Ibrahim E, Khodamoradi K, Delma KS, Iakymenko OA, Kryvenko ON, Arora H, Ramasamy R. Histopathology and Ultrastructural Findings of Fatal COVID-19 Infections on Testis. World J Mens Health. 2021;39(1):65–74. Epub 20201103. doi: 10.5534/wjmh.200170. PubMed PMID: 33151050; PMCID: PMC7752514.

40. Ma X, Guan C, Chen R, Wang Y, Feng S, Wang R, Qu G, Zhao S, Wang F, Wang X, Zhang D, Liu L, Liao A, Yuan S. Pathological and molecular examinations of postmortem testis biopsies reveal SARS-CoV-2 infection in the testis and spermatogenesis damage in COVID-19 patients. Cellular and Molecular Immunology. 2021;18(2):487–9. doi: 10.1038/s41423-020-00604-5.

41. Ardestani Zadeh A, Arab D. COVID-19 and male reproductive system: pathogenic features and possible mechanisms. J Mol Histol. 2021;52(5):869–78. Epub 20210707. doi: 10.1007/s10735-021-10003-3. PubMed PMID: 34232425; PMCID: PMC8260577.

42. Liu X, Chen Y, Tang W, Zhang L, Chen W, Yan Z, Yuan P, Yang M, Kong S, Yan L, Qiao J. Single-cell transcriptome analysis of the novel coronavirus (SARS-CoV-2) associated gene ACE2 expression in normal and non-obstructive azoospermia (NOA) human male testes. Sci China Life Sci. 2020;63(7):1006–15. Epub 20200430. doi: 10.1007/s11427-020-1705-0. PubMed PMID: 32361911; PMCID: PMC7195615.

43. Li X, Lu H, Li F, Zhang Q, Wang T, Qiang L, Yang Q. Impacts of COVID-19 and SARS-CoV-2 on male reproductive function: a systematic review and meta-analysis protocol. BMJ Open. 2022;12(1):e053051. Epub 20220105. doi: 10.1136/bmjopen-2021-053051. PubMed PMID: 34987042.

44. Li C, Ye Z, Zhang AJ, Chan JF, Song W, Liu F, Chen Y, Kwan MY, Lee AC, Zhao Y, Wong BH, Yip CC, Cai JP, Lung DC, Sridhar S, Jin D, Chu H, To KK, Yuen KY. Severe acute respiratory syndrome coronavirus 2 (SARS-CoV-2) infections by intranasal or testicular inoculation induces testicular damage preventable by vaccination in golden Syrian hamsters. Clin Infect Dis. 2022. Epub 20220218. doi: 10.1093/cid/ciac142. PubMed PMID: 35178548.

45. Campos RK, Camargos VN, Azar SR, Haines CA, Eyzaguirre EJ, Rossi SL. SARS-CoV-2 Infects Hamster Testes. Microorganisms. 2021;9(6). Epub 20210617. doi: 10.3390/microorganisms9061318. PubMed PMID: 34204370; PMCID: PMC8235703.

46. Ullah I, Prevost J, Ladinsky MS, Stone H, Lu M, Anand SP, Beaudoin-Bussieres G, Symmes K, Benlarbi M, Ding S, Gasser R, Fink C, Chen Y, Tauzin A, Goyette G, Bourassa C, Medjahed H, Mack M, Chung K, Wilen CB, Dekaban GA, Dikeakos JD, Bruce EA, Kaufmann DE, Stamatatos L, McGuire AT, Richard J, Pazgier M, Bjorkman PJ, Mothes W, Finzi A, Kumar P, Uchil PD. Live imaging of SARS-CoV-2 infection in mice reveals that neutralizing antibodies require Fc function for optimal efficacy. Immunity. 2021;54(9):2143–58 e15. Epub 20210818. doi: 10.1016/j.immuni.2021.08.015. PubMed PMID: 34453881; PMCID: PMC8372518.

47. Pike JFW, Polley EL, Pritchett DY, Lal A, Wynia BA, Roudebush WE, Chosed RJ. Comparative analysis of viral infection outcomes in human seminal fluid from prior viral epidemics and Sars-CoV-2 may offer trends for viral sexual transmissibility and long-term reproductive health implications. Reprod Health. 2021;18(1):123. Epub 20210610. doi: 10.1186/s12978-021-01172-1. PubMed PMID: 34112171; PMCID: PMC8192109.

48. Massarotti C, Garolla A, Maccarini E, Scaruffi P, Stigliani S, Anserini P, Foresta C. SARS-CoV-2 in the semen: Where does it come from? Andrology. 2021;9(1):39–41. Epub 20200728. doi: 10.1111/andr.12839. PubMed PMID: 32533891; PMCID: PMC7323151.

49. Haghpanah A, Masjedi F, Salehipour M, Hosseinpour A, Roozbeh J, Dehghani A. Is COVID-19 a risk factor for progression of benign prostatic hyperplasia and exacerbation of its related symptoms?: a systematic review. Prostate Cancer Prostatic Dis. 2021. Epub 20210518. doi: 10.1038/s41391-021-00388-3. PubMed PMID: 34007019; PMCID: PMC8129694.

50. Stopsack KH, Mucci LA, Antonarakis ES, Nelson PS, Kantoff PW. TMPRSS2 and COVID-19: Serendipity or Opportunity for Intervention? Cancer Discov. 2020;10(6):779–82. Epub 20200410. doi: 10.1158/2159-8290.CD-20-0451. PubMed PMID: 32276929; PMCID: PMC7437472.

51. Gedeborg R, Styrke J, Loeb S, Garmo H, Stattin P. Androgen deprivation therapy and excess mortality in men with prostate cancer during the initial phase of the COVID-19 pandemic. PLoS One. 2021;16(10):e0255966. Epub 20211007. doi: 10.1371/journal.pone.0255966. PubMed PMID: 34618806; PMCID: PMC8496782.

52. Soumarova R, Boday A, Krhutova V, Janotova A, Dvorakova M, Jaluvkova E, Stursa M, Perkova H. Prognostic and predictive molecular biological markers in prostate cancer - significance of expression of genes PCA3 and TMPRSS2. Neoplasma. 2015;62(1):114–8. doi: 10.4149/neo_2015_014. PubMed PMID: 25563374.

53. Bhowmick NA, Oft J, Dorff T, Pal S, Agarwal N, Figlin RA, Posadas EM, Freedland SJ, Gong J. COVID-19 and androgen-targeted therapy for prostate cancer patients. Endocr Relat Cancer. 2020;27(9):R281–R92. doi: 10.1530/ERC-20-0165. PubMed PMID: 32508311; PMCID: PMC7546583.

54. Kazan O, Culpan M, Efiloglu O, Atis G, Yildirim A. The clinical impact of androgen deprivation therapy on SARS-CoV-2 infection rates and disease severity. Turk J Urol. 2021;47(6):495–500. doi: 10.5152/tud.2021.21278. PubMed PMID: 35118968.

55. Cinislioglu AE, Demirdogen SO, Cinislioglu N, Altay MS, Sam E, Akkas F, Tor IH, Aydin HR, Karabulut I, Ozbey I. Variation of Serum PSA Levels in COVID-19 Infected Male Patients with Benign Prostatic Hyperplasia (BPH): A Prospective Cohort Studys. Urology. 2022;159:16–21. Epub 20211006. doi: 10.1016/j.urology.2021.09.016. PubMed PMID: 34626600; PMCID: PMC8493783.

56. Mohamed MS, Moulin TC, Schioth HB. Sex differences in COVID-19: the role of androgens in disease severity and progression. Endocrine. 2021;71(1):3–8. Epub 20201111. doi: 10.1007/s12020-020-02536-6. PubMed PMID: 33179220; PMCID: PMC7657570.

57. Reddy R, Farber N, Kresch E, Seetharam D, Diaz P, Ramasamy R. SARS-CoV-2 in the Prostate: Immunohistochemical and Ultrastructural Studies. The World Journal of Men’s Health. 2021;40. doi: 10.5534/WJMH.210174.

58. Fraga-Silva RA, Costa-Fraga FP, Montecucco F, Sturny M, Faye Y, Mach F, Pelli G, Shenoy V, da Silva RF, Raizada MK, Santos RA, Stergiopulos N. Diminazene protects corpus cavernosum against hypercholesterolemia-induced injury. J Sex Med. 2015;12(2):289–302. Epub 20141120. doi: 10.1111/jsm.12757. PubMed PMID: 25411084.

59. Kresch E, Achua J, Saltzman R, Khodamoradi K, Arora H, Ibrahim E, Kryvenko ON, Wolff Almeida V, Firdaus F, Hare JM, Ramasamy R. COVID-19 Endothelial Dysfunction Can Cause Erectile Dysfunction: Histopathological, Immunohistochemical, and Ultrastructural Study of the Human Penis. World J Mens Health. 2021;39:2287–4208. doi: 10.5534/wjmh.210055.

60. Sansone A, Mollaioli D, Ciocca G, Colonnello E, Limoncin E, Balercia G, Jannini EA. “Mask up to keep it up”: Preliminary evidence of the association between erectile dysfunction and COVID-19. Andrology. 2021;00:1–7. doi: 10.1111/andr.13003.

61. Dal Moro F, Vendramin I, Livi U. The war against the SARS-CoV2 infection: Is it better to fight or mitigate it? Med Hypotheses. 2020;143:110129. Epub 20200722. doi: 10.1016/j.mehy.2020.110129. PubMed PMID: 32721814; PMCID: PMC7373683.

62. Mejri R, Mrad Dali K, Chaker K, Bibi M, Ben Rhouma S, Nouira Y. Venous thrombosis of the pampiniform plexus after coronavirus infection (COVID-19): A case report. Urol Case Rep. 2021;39:101860. Epub 20210927. doi: 10.1016/j.eucr.2021.101860. PubMed PMID: 34603969; PMCID: PMC8475016.

63. Alshoabi SA, Haider KH, Mostafa MA, Hamid AM, Daqqaq TS. An unusual and atypical presentation of the novel coronavirus: A case report and brief review of the literature. J Taibah Univ Med Sci. 2021;16(4):637–42. Epub 20210412. doi: 10.1016/j.jtumed.2021.01.014. PubMed PMID: 33867909; PMCID: PMC8038890.

64. Whiteley MS, Abu-Bakr O, Holdstock JM. Testicular vein thrombosis mimicking epididymo-orchitis after suspected Covid-19 infection. SAGE Open Med Case Rep. 2021;9:2050313X211022425. Epub 20210604. doi: 10.1177/2050313X211022425. PubMed PMID: 34158948; PMCID: PMC8182169.

65. La Marca A, Busani S, Donno V, Guaraldi G, Ligabue G, Girardis M. Testicular pain as an unusual presentation of COVID-19: a brief review of SARS-CoV-2 and the testis. Reprod Biomed Online. 2020;41(5):903–6. Epub 20200723. doi: 10.1016/j.rbmo.2020.07.017. PubMed PMID: 32826162; PMCID: PMC7377719.

66. Li K, Liu X, Huang Y, Liu X, Song Q, Wang R. Evaluation of testicular spermatogenic function by ultrasound elastography in patients with varicocele-associated infertility. Am J Transl Res. 2021;13(8):9136–42. Epub 20210815. PubMed PMID: 34540028; PMCID: PMC8430184.

67. Mali WP, Arndt JW, Coolsaet BL, Kremer J, Oei HY. Haemodynamic aspects of left-sided varicocele and its association with so-called right-sided varicocele. Int J Androl. 1984;7(4):297–308. doi: 10.1111/j.1365-2605.1984.tb00787.x. PubMed PMID: 6096276.

68. Aitken RJ. COVID-19 and human spermatozoa—Potential risks for infertility and sexual transmission? Andrology. 2021;9(1):48–52. doi: 10.1111/andr.12859.

69. Gacci M, Coppi M, Baldi E, Sebastianelli A, Zaccaro C, Morselli S, Pecoraro A, Manera A, Nicoletti R, Liaci A, Bisegna C, Gemma L, Giancane S, Pollini S, Antonelli A, Lagi F, Marchiani S, Dabizzi S, Degl’Innocenti S, Annunziato F, Maggi M, Vignozzi L, Bartoloni A, Rossolini GM, Serni S. Semen impairment and occurrence of SARS-CoV-2 virus in semen after recovery from COVID-19. Human Reproduction. 2021;0(0):1–10. doi: 10.1093/humrep/deab026.

70. Sengupta P, Leisegang K, Agarwal A. The impact of COVID-19 on the male reproductive tract and fertility: A systematic review. Arab J Urol. 2021;19(3):423–36. Epub 20210809. doi: 10.1080/2090598X.2021.1955554. PubMed PMID: 34552795; PMCID: PMC8451696.

71. Li H, Xiao X, Zhang J, Zafar MI, Wu C, Long Y, Lu W, Pan F, Meng T, Zhao K, Zhou L, Shen S, Liu L, Liu Q, Xiong C. Impaired spermatogenesis in COVID-19 patients. EClinicalMedicine. 2020;28. doi: 10.1016/j.eclinm.2020.100604.

72. Delli Muti N, Finocchi F, Tossetta G, Salvio G, Cutini M, Marzioni D, Balercia G. Could SARS-CoV-2 infection affect male fertility and sexuality? APMIS. 2022. Epub 20220203. doi: 10.1111/apm.13210. PubMed PMID: 35114008.

73. Wesselink AK, Hatch EE, Rothman KJ, Wang TR, Willis MD, Yland J, Crowe HM, Geller RJ, Willis SK, Perkins RB, Regan AK, Levinson J, Mikkelsen EM, Wise LA. A prospective cohort study of COVID-19 vaccination, SARS-CoV-2 infection, and fertility. Am J Epidemiol. 2022. Epub 20220120. doi: 10.1093/aje/kwac011. PubMed PMID: 35051292; PMCID: PMC8807200.

74. Xu J, Qi L, Chi X, Yang J, Wei X, Gong E, Peh S, Gu J. Orchitis: A Complication of Severe Acute Respiratory Syndrome (SARS). Biology of Reproduction. 2006;74(2):410–6. doi: 10.1095/BIOLREPROD.105.044776.

75. Delle Fave RF, Polisini G, Giglioni G, Parlavecchio A, Dell’Atti L, Galosi AB. COVID-19 and male fertility: Taking stock of one year after the outbreak began. Archivio Italiano di Urologia e Andrologia. 2021;93(1):115–9. doi: 10.4081/aiua.2021.1.115.

76. Al-Kuraishy HM, Al-Gareeb AI, Faidah H, Alexiou A, Batiha GE. Testosterone in COVID-19: An Adversary Bane or Comrade Boon. Front Cell Infect Microbiol. 2021;11:666987. Epub 20210908. doi: 10.3389/fcimb.2021.666987. PubMed PMID: 34568081; PMCID: PMC8455954.

77. Haghpanah A, Masjedi F, Alborzi S, Hosseinpour A, Dehghani A, Malekmakan L, Roozbeh J. Potential mechanisms of SARS-CoV-2 action on male gonadal function and fertility: Current status and future prospects. Andrologia. 2021;53(1):e13883-e. doi: 10.1111/and.13883.

78. Enikeev D, Taratkin M, Morozov A, Petov V, Korolev D, Shpikina A, Spivak L, Kharlamova S, Shchedrina I, Mestnikov O, Fiev D, Ganzha T, Geladze M, Mambetova A, Kogan E, Zharkov N, Demyashkin G, Shariat SF, Glybochko P. Prospective two-arm study of the testicular function in patients with COVID-19. Andrology. 2022. Epub 20220206. doi: 10.1111/andr.13159. PubMed PMID: 35124885.

